# Within- and between-host dynamics of highly pathogenic avian influenza in domestic birds from Pennsylvania farms and live bird markets

**DOI:** 10.64898/2026.07.17.739193

**Authors:** Anna S Jaeger, Elena Cruz-Adames, Stephen D Shank, Irina Chupikova, Katie Kumta, Lisa A Murphy, Eman Anis, Louise H Moncla

**Affiliations:** Department of Pathobiology, School of Veterinary Medicine, University of Pennsylvania

## Abstract

Since late 2021, highly pathogenic H5 avian influenza viruses have spread across the Americas, resulting in hundreds of outbreaks in agriculture and 71 human spillovers. In the United States, human infections overwhelmingly derive from exposure to infected agricultural animals. However, little is known about how these viruses evolve during outbreaks in agriculture, or how they spread locally, key information for understanding human infection risk. We produced high-quality deep sequence data from commercial and live bird market (LBM) outbreaks in Pennsylvania from 2023-2025. We uncover repeated spillovers from wild to domestic birds, and evidence of high circulating diversity in LBMs that likely moved across state lines. Samples collected from the same outbreak shared a significant fraction of viral variation, suggesting high circulating diversity and a strong signal of transmission linkage. Samples from poultry collectively harbored 9 known adaptive variants, including PB2 D701N, suggesting that rare, adaptive variants may circulate in settings where human exposure risk is high. Our data suggest that outbreaks in poultry pose opportunities for subsequent viral evolution, supporting continued efforts to stamp out transmission. We also find that LBMs experience outbreak risk from domestic and wild sources and promote viral mixing between species, suggesting that enhanced tracking and biosecurity may be useful in these settings. Finally, our data suggest the potential to use within-sample variation for enhanced outbreak linkage within these outbreak settings.

Since 2021, highly pathogenic avian influenza viruses have caused hundreds of agricultural outbreaks and 71 human spillovers. Human infections overwhelmingly derive from infected agricultural animals, but little is known about how these viruses evolve and spread locally. We generated deep sequence data from commercial and live bird market (LBM) outbreaks in Pennsylvania, USA from 2023-2025. We uncover repeated spillovers from wild to domestic birds, and high circulating diversity and multispecies mixing in LBMs, resulting in cross-state transmission. Samples from the same outbreak shared substantial viral variation, suggesting high circulating diversity and potential to use viral variation for enhanced transmission linkage. We identified 9 known adaptive variants in poultry, including PB2 D701N, suggesting that rare, adaptive variants may circulate in agricultural settings. Our data suggest that outbreaks in poultry pose opportunities for viral evolution, and that LBMs are high risk settings that may benefit from enhanced monitoring.

## INTRODUCTION

In late 2021, highly pathogenic H5N1 viruses were introduced into the Americas and subsequently spread across the continent ^1,2^. Unlike past outbreaks that were predominantly spread among commercial poultry, the epizootic since 2021 has been driven by wild migratory birds ^3^, making control near impossible. Introductions into agricultural settings have been repeated and ongoing, with dozens to hundreds of introductions leading to the culling of over 200 million domestic birds ^4–6^. Outbreaks in poultry have led to 24 human infections, while additional spillovers have caused outbreaks in dairy cattle and peridomestic species, posing ongoing challenges for animal health and human exposure risk.

From 2021-2026, 71 human spillovers of H5N1 have occurred in the United States. Human infection with H5N1 viruses is primarily mediated by direct interaction with infected animals, with the highest exposure risk during agricultural outbreaks. However, how outbreaks unfold locally, and the capacity for these viruses to evolve during these outbreaks are not well characterized. Analyses of transmission to date have focused on large geographic scales ^1–3^, with limited details on farm types, host species, or local geographic data, all of which likely vary locally to impact risk. Similarly, studies of viral adaptation and evolution have primarily focused on detailing continent-wide patterns of reassortment and genotype formation, identifying consensus-level adaptive mutations in cattle and other emergent lineages, and laboratory studies of putative adaptive mutations ^1,7–10^. Understanding local spread is key for formulating effective outbreak response strategies and biosecurity updates, which are generally developed and managed at the state and local level. Because human infection arises from direct interaction with sick animals, understanding the capacity of these viruses to generate and propagate new mutations is essential for understanding human infection risk in these settings.

Influenza viruses make mutations during the course of infection, resulting in viral diversity within infected individuals. Recent advances in genomic epidemiology have shown that within-host viral variation is shared between donor-recipient pairs when circulating diversity is high and when a large number of virions are transmitted between hosts (e.g., a “wide transmission bottleneck”)(e.g., viruses like HIV, HCV) ^11–14^. In these cases, within-host diversity data could potentially enhance local-scale transmission inference and provide data on the strength of selection. For respiratory viruses like influenza and SARS-CoV-2, variant transmission depends on transmission route ^15,16^, with airborne transmission among humans imposing a strict bottleneck that purges within-host diversity ^17–19^. For avian influenza viruses, outbreak transmission may involve multiple modes of transmission within dense and confined spaces, potentially impacting variant transmission. To date, studies of avian influenza evolution have focused on measuring evolution during experimental infections in animal models^20–22^, and the detection of consensus-level adaptive variants at the population scale ^7–9^, with very few datasets suitable for studying how these viruses adapt during natural outbreaks.

Pennsylvania supports a complex agricultural system that has been heavily impacted by HPAI. Since 2021, over 15 million domestic birds have been affected in Pennsylvania alone, deriving from outbreaks in commercial poultry, backyard flocks, and live bird markets ^23^. Pennsylvania is a major egg producer ^24^, and serves as the primary producer and key distributor for the Northeast live bird market (LBM) system, which connects Pennsylvania, New York and New Jersey. The Northeast LBM system is a complex network that has been heavily impacted by HPAI and accounts for 86.7% of LBM outbreaks in the US since 2022 ^25,26^. LBMs are at particular risk for HPAI due to high levels of bird movement, high diversity of bird species, and intensive mixing from multiple sources ^27^, making viral tracking and outbreak control challenging. Pennsylvania therefore provides a valuable case study for clarifying the degree to which outbreaks among commercial, backyard, and live bird markets are related, and quantifying how diversity is propagated within these settings where humans and animals interact.

We partnered with the Pennsylvania Animal Diagnostic Laboratory System to sequence 46 HPAI positive samples collected from domestic Anseriforme and Galliforme species sampled in Pennsylvania commercial and LBM settings from 2023-2025, along with 39 samples from wild birds. Our data uncover high circulating diversity within commercial farms and LBMs, and the potential for host-adaptive mutations to be present in samples from agricultural outbreaks. We find that live bird market outbreaks experience outbreak risk from wild and domestic sources, support viral mixing among species, and support outbreaks that span multiple states. Finally, we find that within-sample variation is frequently shared among samples from the same outbreak, supporting the potential utility of within-sample diversity data to link outbreaks together. Collectively, our data suggest that agricultural outbreaks represent opportunities for subsequent viral evolution, supporting continued efforts to monitor and eliminate transmission in these settings.

## RESULTS

### Sample information and controls

From 2022-2025, Pennsylvania reported 135 outbreaks among domestic birds, including 60 from commercial flocks, 73 from backyard birds, and 2 from live bird markets ^25^. The Pennsylvania Animal Diagnostics Laboratory System (PADLS) performs diagnostic testing for HPAI suspected cases, and sends non-negatives to the US Department of Agriculture for confirmation. Between 2023 and 2025, 46 samples from domestic birds and 39 samples from wild birds had sufficient RNA for sequencing. These samples span various host species, production types, and sample collection and pooling strategies, which vary across species according to USDA collection procedures ^28^. For domestic birds, gallinaceous poultry specimens consist of up to 11 pooled tracheal (TR) or oropharyngeal (OR) swabs, domestic waterfowl include up to 5 pooled cloacal (CL) swabs, and live bird market (LBM) pools vary due to opportunistic sampling of animals present at market ^28^. Wild bird samples generally represent a single infected animal. Given the variability of pooling methods (Table 1), we opted to use consensus genomes from wild birds purely as contextual data for domestic outbreak reconstructions. For domestic bird samples, we refer to sub-consensus variants as within-sample (as opposed to within-host) variation (iSNVs), and interpret this variation as a proxy for circulating diversity within that pool of samples.

**Table 1:**
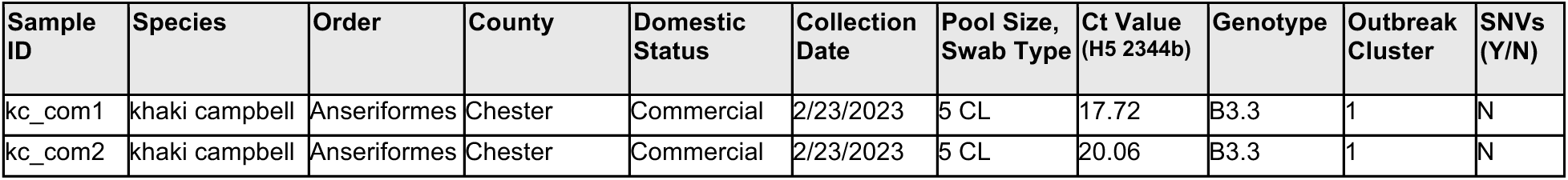
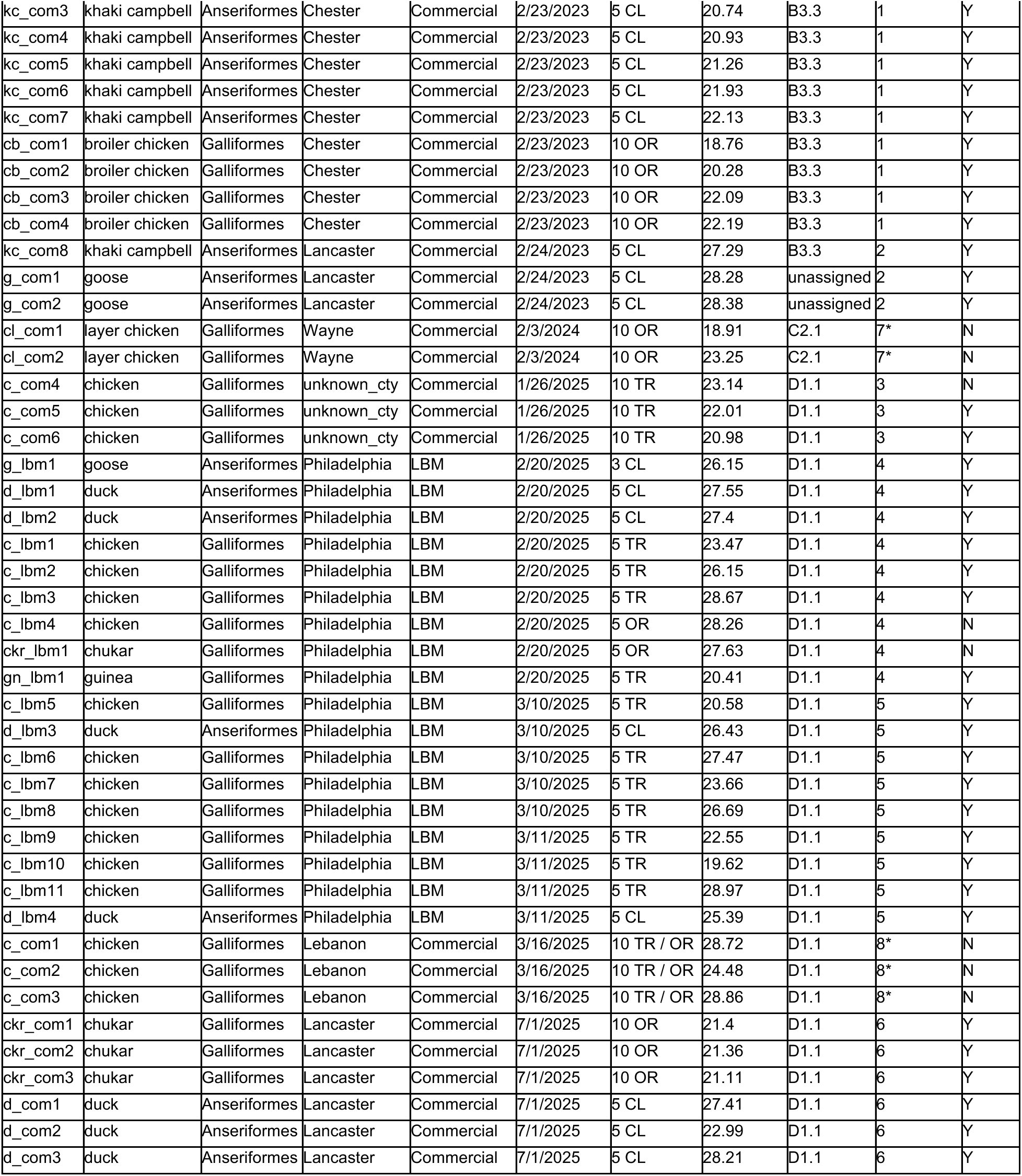
Sample information. CL, cloacal; TR, tracheal, OR, oropharyngeal. Genotype “unassigned” indicates insufficient coverage across gene segments to determine genotype. Outbreak clusters with (*) indicate clusters with no within-sample variants.

Viral deep sequencing data are susceptible to multiple sources of error from PCR amplification, laboratory preparation, and bioinformatic analysis ^18,29,30^. To control for these errors, we developed a sequencing pipeline to produce high-coverage sequencing depth, and sequenced all samples in technical duplicate (see Methods for details). In line with prior work, we find a positive correlation between iSNVs and sample Ct when using iSNVs detected in only one replicate (Figure 1A, slope = 1.202, R^2^=0.06), but not when using variants found in both replicates (Figure 1A, R^2^ = 0.134, slope = −0.582). Variants detected in both technical replicates had generally concordant frequencies (Figure 1B, R^2^ = 0.77, slope = 0.88), supporting the use of technical replicates to control for iSNV variability. For all subsequent analyses, we report iSNVs that were detected in at least 2% of sequencing reads, in both technical replicates, with at least 100x coverage (see Methods for more details). In total, 46 domestic samples (28 commercial farm samples, 18 LBM) matched these criteria and were used for subsequent analysis.

**Figure 1.**
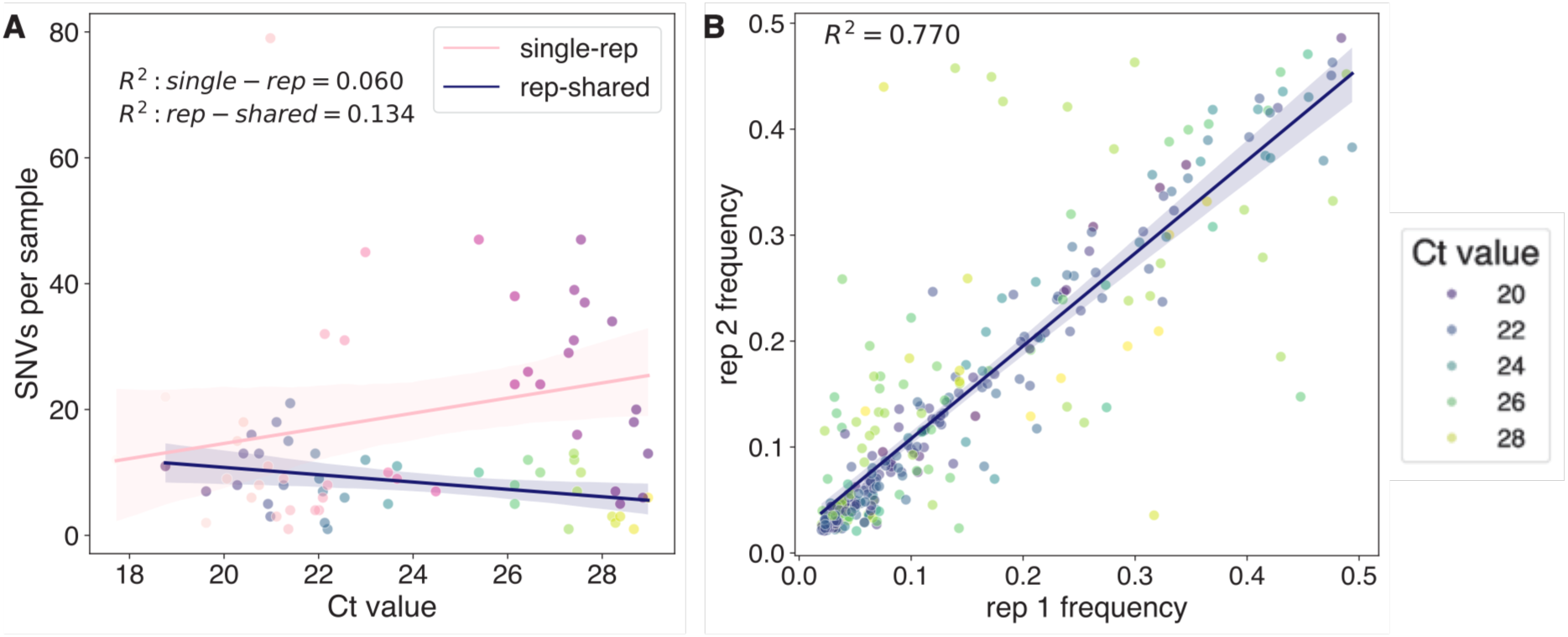
Within-sample variants are reproducible when shared between technical replicates. **A)** The number of SNVs per sample that are shared between technical replicates are slightly negatively correlated with Ct-value. The number of SNVs in only one replicate per sample is positively correlated with Ct-value. **B)** For within-sample minority variants present in both replicates, frequencies in replicate 1 are plotted on the x-axis and frequencies in replicate 2 are plotted on the y-axis. Points represent individual samples colored by Ct-value gradient (single-rep, pink-purple; rep-shared, blue-green). Lines are fitted linear regression lines to single-rep values (pink) and rep-shared values (blue).

### Pennsylvania outbreaks comprise multiple, independent outbreak clusters

Pennsylvania has been heavily impacted by HPAI, with outbreaks reported across commercial poultry, backyard birds, and live bird markets ^31^. To investigate how Pennsylvania outbreaks relate to circulating diversity across the continent, we combined all of our wild and domestic bird consensus sequences with a large, contextual dataset comprised of every publicly available sequence from Pennsylvania (n=233-243 sequences per gene) and ∼6,000 additional North American sequences sampled throughout the panzootic (n=6,593-6,670 sequences per gene) (Supplementary Figure 1). We classified sequences as “Pennsylvania”, “border state”, or “non-border state”, built maximum likelihood trees for each gene segment, and inferred introductions into Pennsylvania using a discrete trait model. From 2022-2025, we infer an average of 73 independent introductions into Pennsylvania (range 68-77 across gene trees, Supplementary Figure 2), including 12-21 that led to transmission within domestic birds (Figure 2A), suggesting a high degree of viral introductions into the state.

**Figure 2.**
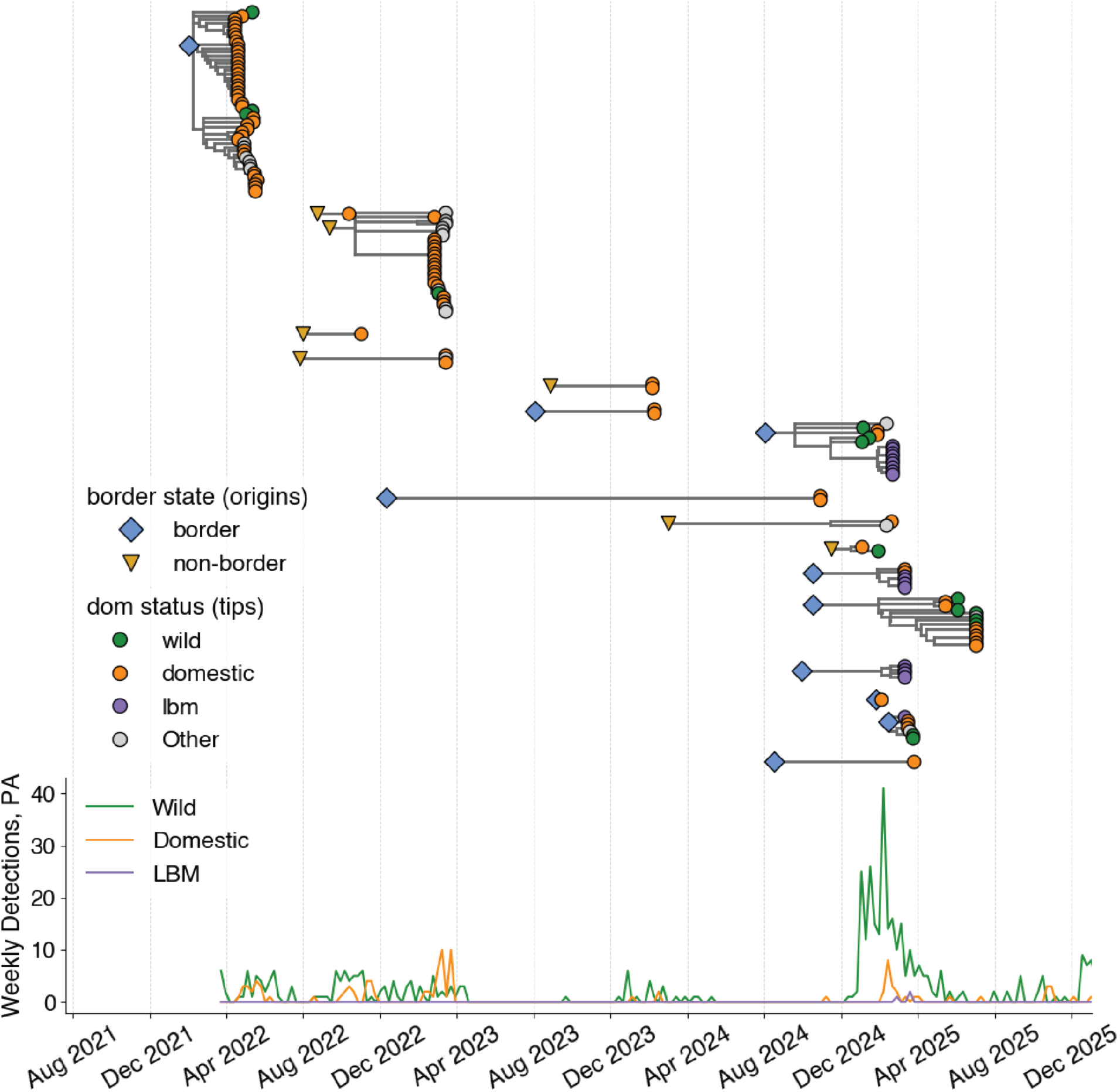
There were many independent introductions into domestic birds in Pennsylvania. Each introduction into Pennsylvania that included domestic infections was inferred from the maximum likelihood time-resolved phylogeny and plotted independently. Large colored points represent the inferred source (border state: diamond, light blue; non-border state: triangle, yellow) that seeded each Pennsylvania subtree. Tips represented by circles are sampled viruses from Pennsylvania colored by domestic status (wild, green; domestic, orange; live bird market (LBM), purple; other/unknown, gray. Subtrees shown here are for the HA phylogeny. Weekly detections from Pennsylvania are shown over time on the shared x-axis for wild, domestic, and LBM birds.

We next grouped our domestic samples into 8 unique outbreak clusters, defined as samples collected from the same Pennsylvania county within the same week. In 2023, our dataset includes two outbreak clusters among commercial poultry reported of the B3.3 genotype (clusters 1 and 2). In 2025, our dataset includes 4 clusters of the D1.1 genotype: cluster 3 represents a commercial poultry outbreak from early 2025; clusters 4 and 5 represent live bird market outbreaks reported in late February and mid March of 2025; and cluster 6 represents a gamebird farm outbreak in July 2025. Two additional clusters were designated, but harbored no within-sample variants and were thus excluded (for more details on outbreak clusters, see Table 1).

Samples from these epidemiologically defined clusters tended to group together on the phylogeny, but separately from each other, suggesting independent origins. To further resolve the relationships between the 4 outbreaks sampled in 2025, we combined all publicly available D1.1 sequences, concatenated genomes, and built a maximum likelihood phylogeny (n=661 full genomes) (Supplementary Figure 3). Concatenated genome trees provide enhanced resolution for genomic inference, particularly when individual genes are identical ^32^. Within the concatenated full genome tree, sequences from each cluster nest within distinct clades, consistent with independent introductions into each outbreak setting (Supplementary Figure 3). These results suggest that the sampled outbreaks in this dataset were not connected to each other, and were likely seeded independently from unsampled sources.

### Pennsylvania outbreaks in live bird markets were seeded independently

Within our dataset, clusters 4 and 5 represent live bird market outbreaks that occurred in the same county, three weeks apart. Given the close location and timing of these outbreaks, we next sought to understand how these outbreaks were related to other viruses sampled in the Northeast. LBMs are susceptible to multiple routes of exposure to HPAI, including from wild birds that are present in or near markets, or from domestic birds that co-mingle during movement through the LBM network, including at holding facilities or the markets themselves. Given the close timing of these outbreaks, they could have been sourced from undetected transmission within the markets themselves, from a common source of infection (e.g., an infected supplier), or from independent introductions from wild birds. Using the collated D1.1 dataset from above, we further classified sequences based on US state of sampling and on domestic/wild status, and modelled transmission using a discrete trait model. These data reveal that the February 2025 LBM outbreak nests most closely with viruses sampled from wild birds. However, the branch connecting these viruses is too distant to confirm a direct link, implying an outbreak source that wasn’t sampled. In contrast, the March 2025 LBM outbreak falls directly within the diversity of viruses sampled from domestic birds in New York at the same time.

Viruses sampled from birds in the Pennsylvania LBM are very genetically similar to those sampled in New York domestic birds, differing by as few as 2 nucleotide mutations in some cases, suggesting very close ancestry. While metadata on sequences from New York in that time period are limited, case detection data specifies whether cases were sampled from commercial or LBM settings. Within the time period of February and March 2025, 20 out of 28 reported domestic outbreaks in New York and New Jersey were from LBMs, suggesting that most domestic infections derived from LBMs (Figure 3). Given this heavy skewing and the known connectivity between the Pennsylvania and New York live bird market systems, we speculate that this cluster represents transmission between LBMs in New York and Pennsylvania. Bayesian phylogeographic analysis infers well-supported transition rates from domestic birds in New York to those in Pennsylvania (Supplementary Table 1), and infers New York domestic birds as the source of clusters of domestic bird infections in New York, New Jersey, and Pennsylvania during this time period (Figure 3). Together, these data show that the Pennsylvania LBM outbreaks in the same county and time had entirely distinct origins, suggesting that LBMs are at risk from myriad domestic and wild sources. Further, our data suggest that viruses from Pennsylvania LBMs are closely connected to those from New York domestic birds, implying that transmission between domestic production systems in these states may have occurred.

**Figure 3.**
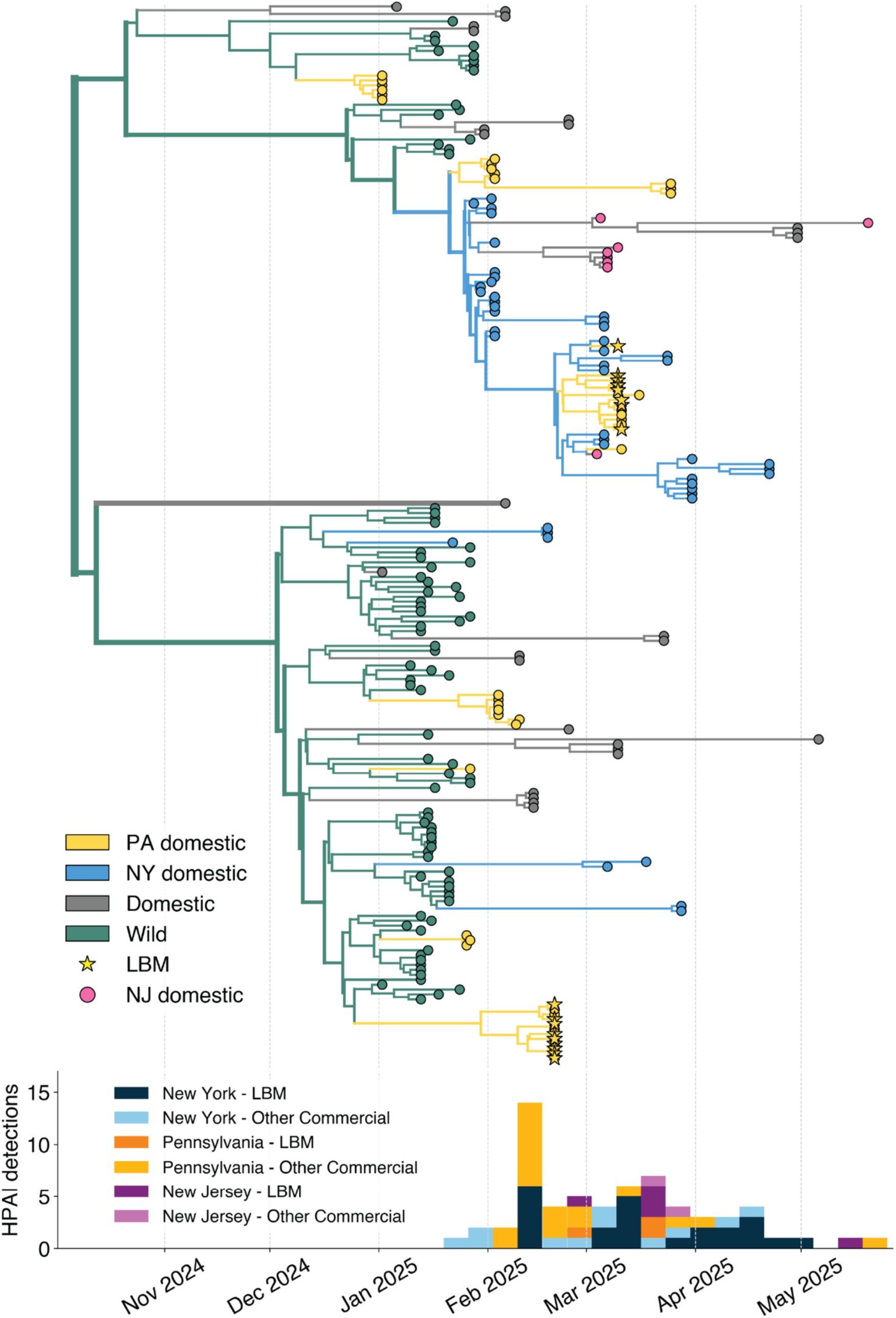
There were multiple introductions into the Pennsylvania live bird market system. A Bayesian phylogenetic reconstruction of transmission between 215 full-genome D1.1 sequences from PA (yellow) and NY (blue) domestic birds, other domestic birds (gray), and wild birds (teal). Sequences from NJ domestic infections are colored in pink, but are not a discrete trait included in the model. The maximum-clade credibility tree is shown here with tips and branches colored by these discrete traits. Weekly HPAI detections from NY, PA, and NJ LBM and commercial production are plotted on the shared x-axis.

### Within-sample diversity does not vary by host or production type, but is higher in D1.1 samples

Our data suggest that HPAI outbreaks in Pennsylvania were repeatedly, independently seeded into commercial and LBM settings, and that LBM outbreaks support complex outbreak dynamics that may move between states. These outbreaks represent key interfaces at which human exposure risk is high, and provide potential opportunities for novel diversity to be generated and propagated. To quantify how much viral variation circulates within these settings, we used our replicate deep sequence data to identify all within-sample variants (iSNVs, representing variants present at ≤50% frequency) in each domestic sample. Because our samples represent pooled swabs, within-sample diversity could reflect a combination of within and between host variation, thus providing a useful proxy for viral diversity within that pool of samples. We identified a total of 307 iSNVs (188 synonymous and 119 nonsynonymous) across 36 high coverage samples spanning all influenza coding regions (Figure 4A). Samples had 8.5 variants (standard deviation ± 5) on average, with variants found across a range of frequencies (Figure 4B). We found no significant difference in the mean number of iSNVs between samples from Anseriformes and Galliformes, or between LBM and commercial farm samples (Figure 4C-E). However, samples from D1.1 genotype viruses had a significantly higher mean number of iSNVs than B3.3 samples (D1.1 mean: 0.798 SNVs/1kb coverage; B3.3 mean: 0.511 SNVs/1kb coverage, t-test, *p* = 0.0239)(Figure 4D). Genotype was the strongest predictor of the number of iSNVs per 1 kb, (beta = 0.420, 0.095-0.745, 95% CI, *p*=0.01), although the model was only modestly predictive overall (adjusted R^2^ = 0.157) (Figure 4F). These data suggest that diversity was similar in samples collected in different species and production types, with some signal of increased diversity in D1.1 genotype infections.

**Figure 4.**
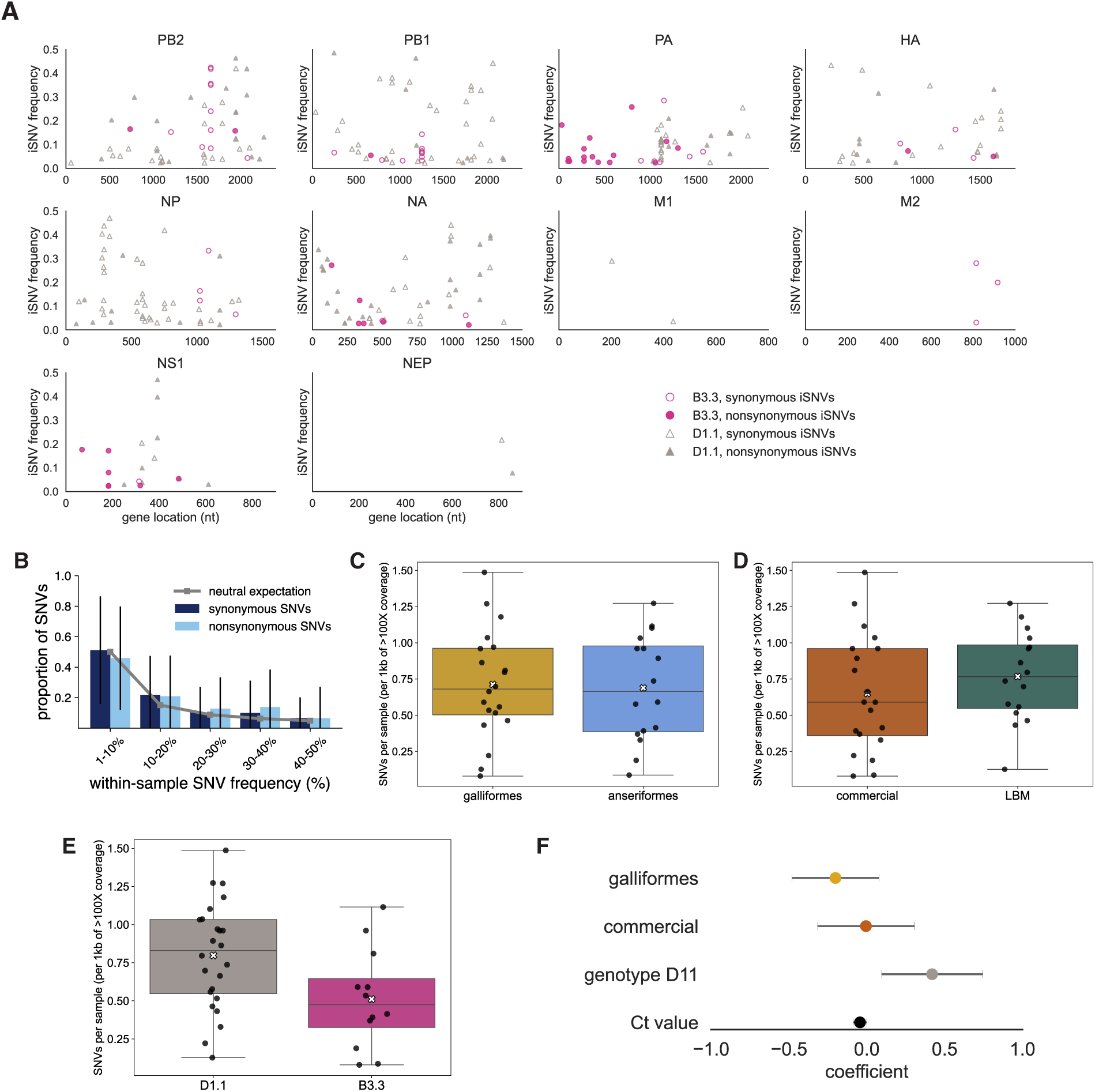
Within-sample variation is low across host types, domestic operations, and genotypes. A) Within-sample variants present in both technical replicates at ≥2% of sequencing reads in all samples across all genes. Each point represents an individual single nucleotide variant (iSNV) present in a sample colored by genotype (B3.3, magenta; D1.1, gray). The x-axis represents the nucleotide position of the SNV for each gene, and the y-axis represents its average frequency within-sample. Open symbols represent synonymous SNVs, closed symbols represent nonsynonymous SNVs. B) The proportion of synonymous (dark blue) and nonsynonymous (light blue) SNVs for each sample was calculated for each frequency bin on the x-axis. Bars represent the mean proportion of variants in each bin with standard deviation represented by black vertical lines. The gray line indicates the expected mean proportion of variants in each bin under a neutral model. The mean number of SNVs per sample per 1kb of ≥ 100X sequencing coverage was compared between Anseriformes and Galliformes (C), commercial farm and LBMs (D), and B3.3 and D1.1 genotypes (E). Mean values are shown by white Xs, individual sample values are shown by black points. F) Only D1.1 genotype had a significant positive effect on the number of SNVs per sample in a linear regression model.

### Putative mammal-adaptive mutations are rare, but present in domestic bird samples

Influenza viruses are thought to require adaptive mutations to infect mammals, which frequently arise in the polymerase complex and surface receptor genes ^33,34^. Given the opportunity for human exposure during domestic bird outbreaks, we compiled a reference database of previously annotated functional sites, and screened our within-sample variants against these sites ^35–37^. We identified 50 iSNVs in our dataset present at 35 unique functional variant sites, 21 of which were nonsynonymous (Figure 5A). iSNVs at functional sites were more frequent in Galliformes (29/50) than Anseriformes (21/50), and in LBM samples (35) compared to commercial farm samples (15). Of the 21 nonsynonymous variants, 9 encoded for the specific amino acid mutation that is cited as adaptive (Table 2). These mutations include HA D171N present in a LBM Anseriformes infection, which is located in an antibody binding site that has been identified in cattle in the US, sheep in the UK, and in South American HPAI isolates ^9,38,39^. PB2 K339T was present in two LBM samples and has been shown to increase H5N1 replication in mammals in laboratory studies ^40^. Most notably, PB2 D701N was present in a chicken sample from a live bird market at 4.8% frequency. PB2 D701N has been repeatedly linked to increased replication efficiency in mammals. This mutation has arisen multiple independent times across the North American H5N1 phylogeny ^41–43^, and was recently shown to enhance transmission of D1.1 viruses in ferret studies ^44^. The chicken sample in our dataset was not phylogenetically linked to any other available sequence that had a PB2 D701N mutation at consensus, suggesting that this mutation was likely introduced from an unsampled source, or independently generated during the outbreak (Figure 5B). These data suggest that while rare, putative mammal-adaptive mutations are tolerated and may be present within agricultural settings where humans and birds interact. These data suggest that domestic bird outbreaks provide evolutionary opportunities for viral adaptation, and support continued efforts to stamp out transmission.

**Figure 5.**
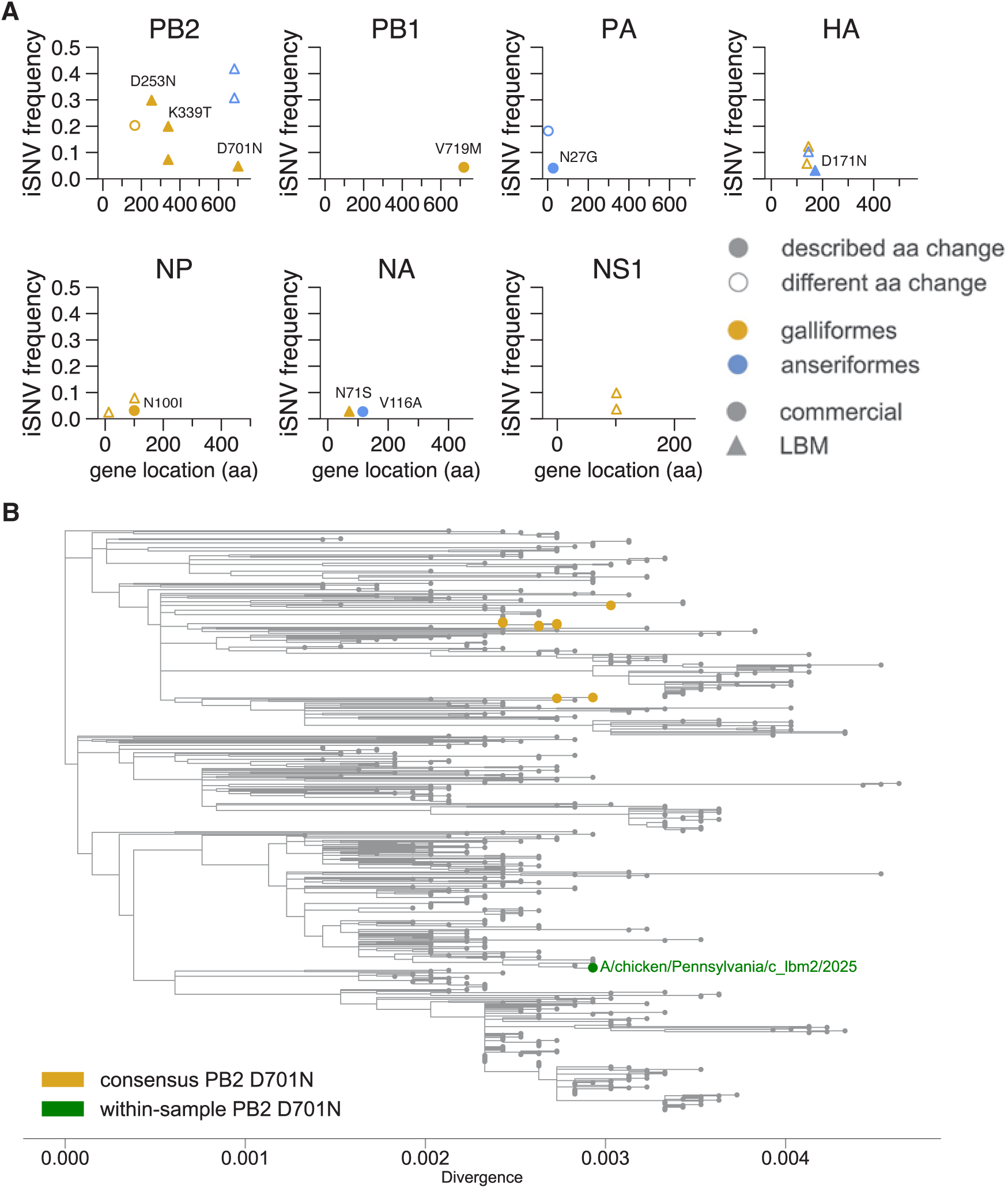
Functional SNVs are present within-sample in domestic infections. A) 21 nonsynonymous mutations were identified within-sample at previously annotated functional sites. Points are colored by host order (yellow, Galliformes; blue, Anseriformes) and shapes indicate domestic status (circle, commercial; triangle, LBM). Open symbols indicate a different amino acid substitution than has been previously cited in literature. Closed symbols indicate the identical amino acid substitution that has been previously identified, and the specific position and change is labeled. The functional iSNV frequency is indicated on the y-axis and the amino acid position in each gene is shown on the x-axis. B) A maximum-likelihood divergence phylogeny of D1.1 sequences show that the infection containing PB2 D701N within-sample (green) is phylogenetically distinct from any sequences with this mutation at consensus (yellow).

**Figure 6.**
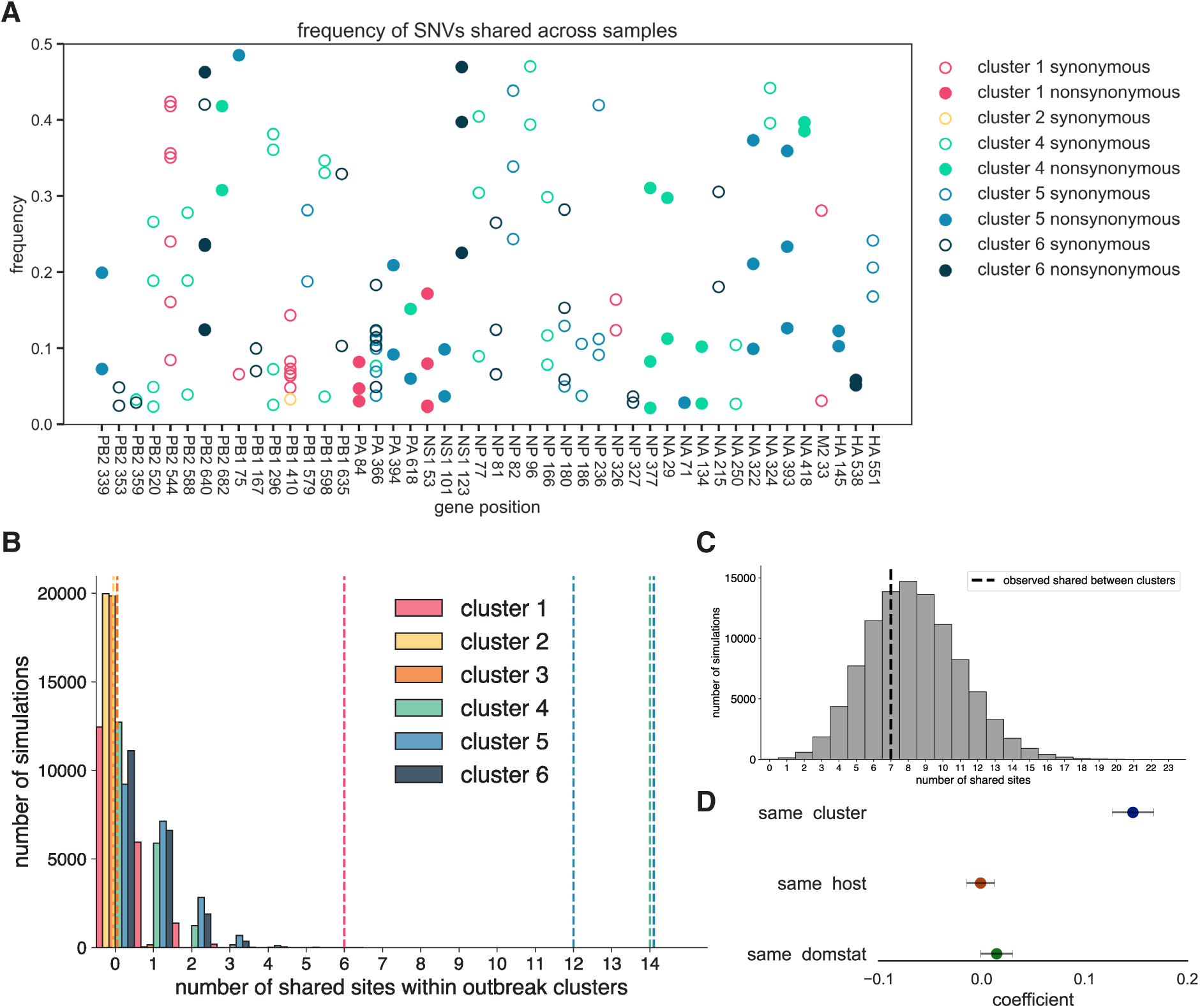
Within-sample variants are shared among infections within outbreak clusters. A) All amino acid sites that had SNVs in at least 2 samples are shown with their gene and amino acid position plotted on the x-axis. Open symbols indicate synonymous SNVS and closed symbols indicate nonsynonymous SNVs. Points are colored by their outbreak cluster, which is grouped by date and county. Frequency is shown on the y-axis. B) A permutation test was performed to determine if the level of within-cluster sharing was more than expected by chance. The number of sites shared by at least two samples within an outbreak cluster is plotted on the x-axis and the bar height indicates the number of simulations with that number of shared sites (20000 total simulations per cluster). Dotted lines represent the observed number of shared sites in each outbreak cluster in the data. C) A second permutation test determined if the level of sharing between clusters was more than expected by chance. The number of shared sites shared by at least two samples in different clusters was plotted on the x-axis and the bar height indicates the number of simulations with that number of shared sites (100,000 simulations). The dotted line (black) represents the observed number of shared sites between clusters in the data. D) We assessed the impact of samples being from the same outbreak cluster, same host order, and same domestic status on the proportion of SNVs shared between each pair of samples in the dataset using a linear regression. The mean of each estimated coefficient in the linear regression model containing all three predictor variables are plotted on the x-axis with the error bars indicating the 95% confidence intervals.

**Table 2:** Within-sample functional mutation information.

| Sample | Gene | Nt position | Coding region change | Avg. frequency | Production type | Order | Genotype | Description |
| --- | --- | --- | --- | --- | --- | --- | --- | --- |
| c_lbm5 | PB2 | 1043 | Lys339Thr | 0.07 | LBM | Galliformes | D1.1 | Increased mammalian virulence, H5N1 <sup>40</sup> |
| c_lbm8 | PB2 | 1043 | Lys339Thr | 0.20 | LBM | Galliformes | D1.1 | Increased mammalian virulence, H5N1 <sup>40</sup> |
| c_lbm2 | PB2 | 2128 | Asp701Asn | 0.05 | LBM | Galliformes | D1.1 | Increased mammalian replication, H5N1 <sup>41,42,44</sup> |
| kc_com3 | PA | 104 | Asp27Gly | 0.04 | commercial | Anseriformes | B3.3 | Reduced antiviral susceptibility <sup>45</sup>; mouse adaptation, H1N1 <sup>46</sup> |
| d_lbm3 | HA | 539 | Asp171Asn | 0.03 | LBM | Anseriformes | D1.1 | Antibody binding site-potential immune evasion and mammalian adaptation, H5N1 <sup>9,38,39,47</sup> |
| ckr_com1 | NP | 344 | Arg100Ile | 0.03 | commercial | Galliformes | D1.1 | Increased virulence in mice, H1N1 <sup>48</sup> |
| c_lbm7 | NA | 232 | Asn71Ser | 0.03 | LBM | Galliformes | D1.1 | Host adaptation- H5N1 cat infections <sup>49,50</sup> |
| kc_com6 | NA | 367 | Val116Ala | 0.03 | commercial | Anseriformes | B3.3 | Reduced antiviral susceptibility, H5N1, H3N2, H1N1 <sup>51-53</sup> |
| c_lbm9 | PB2 | 253 | Asp253Asn | 0.30 | LBM | Galliformes | D1.1 | Increased mammalian replication, H9N2 <sup>54,55</sup> |
| ckr_com1 | PB1 | 2179 | Val719Met | 0.04 | commercial | Galliformes | D1.1 | Increased virulence in mice with PB2 627K in H7N9 <sup>56</sup> |

### Within-sample variation is shared among outbreak samples

Prior studies of influenza within-host diversity have found that infected individuals harbor very low within-host diversity and that variants are rarely shared between individuals ^17,18,57^. This phenomenon is true even among putative household transmission pairs, and likely arises from a tight bottleneck that purges within-host diversity during transmission. In our dataset, several variants were detected at mid to high frequencies (30-50%, Figure 4a), and we observe a smaller fraction of variants at very low frequencies (<10%) than prior studies (Figure 4b). We also noted several variants that were found in multiple samples, including some variants that were present in up to 11 unique pools. While this phenomenon may arise from the pooling process, the unexpectedly high level of diversity suggests that shared variants could potentially be useful for linking epidemiologically related samples. We thus next sought to determine whether within-sample variation was shared among samples from the same outbreak.

We identified 47 total polymorphic sites in which a variant was detected in two or more samples (Figure 5A). Among these 47 shared sites, 19 (40%) were nonsynonymous, while 28 (60%) were synonymous. Compared to the proportion of nonsynonymous and synonymous sites in the H5N1 genome (∼74% nonsynonymous, ∼26% synonymous), these shared sites in our dataset reflect a significant excess of shared synonymous variants (Chi-square with Yate’s correction, *p* < 0.0001)^58^. These data suggest that these shared variants more likely arise from epidemiologic linkage, rather than from convergent evolution.

Next, we labelled each variant with information on its epidemiologically-defined outbreak cluster (samples collected from the same county in the same week). 43 out of 47 shared sites were shared exclusively between samples from the same cluster. To determine whether within-cluster variant sharing was statistically significant, we simulated the same number of mutations as observed in the actual data, but randomized the sites across the genome. We simulated this process 20,000 times, and counted the number of shared sites to generate null distributions for each cluster (see Methods for details). We then compared the observed shared variants for each cluster to its null distribution (Figure 5B). Clusters 2 and 3 included 2-3 total variants, with none shared, and were not significant. However, clusters 1, 4, 5, and 6 shared significantly more variants than expected by chance (*p* < 0.0001). Clusters 4 and 5 (representing the LBM outbreaks) harbored the highest proportion of shared variants, implying a high degree of diversity and mixing within these settings. In contrast, the number of sites shared between clusters did not differ from the null (n=7) (Figure 5C), suggesting that variants are rarely shared among samples from different epidemiologic events. In a linear regression model, being in the same outbreak cluster was the best predictor of sharing variants, while being from the same domestic setting or host order had no effect (Figure 5D). These results suggest that both commercial and LBM outbreaks support high levels of circulating diversity that is frequently shared among samples collected from the same outbreak. Pooled outbreak samples thus retain genetic signatures of epidemiologic linkage, suggesting potential for the use of these data to enhance transmission linkage when sampling depth is high.

### Commercial and LBM outbreaks show distinct mixing and host-specific patterns

Given the high degree of circulating diversity and shared variation, we next wanted to determine whether combining within-sample variation with consensus data would provide additional information on outbreak spread and mixing. We used consensus sequences from our outbreak clusters with significant variant sharing (clusters 1, 4, 5, and 6), along with publicly available sequences from the same genotype and time period (B3.3 and D1.1) to build concatenated, full genome divergence trees (Supplementary Figure 4A-D). We also examined shared variants within each setting, subsetted by host (Supplementary Figure 4E-F). These data reveal differences in outbreak mixing in the commercial and LBM outbreaks in our dataset. In the B3.3 phylogeny (cluster 1, commercial outbreak), our samples form a distinct cluster with many of our sequences identical at the consensus level. Within this cluster, sequences from duck samples cluster more basal in the tree, falling upstream of infections sampled in chickens. This structure is also mildly apparent in cluster 5 of the D1.1 tree (cluster 5, commercial outbreak in July 2025) (Supplementary Figure 4B). This pattern is consistent with a model in which infections in ducks occurred earlier, with subsequent transmission to chickens. Within these two commercial clusters, variants were mostly shared among samples from the same hosts. Among the 17 shared polymorphic sites in commercial samples, 5 were detected in multiple hosts, while 12 were host specific. These data imply some degree of host separation in viral diversity, with viral infections in ducks and chickens proceeding independently.

In contrast, clusters involving LBM samples are more mixed. LBM outbreak samples (clusters 4 and 5) form larger, more diverse trees, with high apparent mixing between samples from Anseriformes and Galliformes. In particular, the March cluster involves substantial mixing between species and between samples from New York and Pennsylvania. LBM clusters also showed far higher rates of within-sample variation that was shared between species, with 21 out of 28 shared variants shared between species. These data align with the known diversity of hosts present at LBMs, and suggest that these markets may promote greater mixing of viral diversity among species. Together, our data highlight that genomic data across scales can reveal differences in outbreak dynamics between production settings. Consensus and within-sample data suggest that these commercial outbreaks supported lower overall viral diversity, greater separation in viral variation between species, and duck infections that were basal to infections in chickens. In contrast, the two LBM outbreaks in our data exhibited higher circulating diversity that was connected across geographic locations and host species, suggesting the potential for these markets to mix viral variation from disparate locations and hosts. The outbreak clusters in our dataset represent only a small subset of total outbreaks in the state, and characterization of more outbreaks is necessary to determine whether these patterns are broadly true. However, these data suggest that LBMs represent high risk interfaces, at which viruses can mix across states and species, and propagate viral diversity in settings where human exposure risk is high.

## DISCUSSION

Agricultural outbreaks represent key interfaces for human infection. Since 2021, outbreaks of H5N1 among domestic birds have been persistent and challenging to control, posing challenges for agriculture and human exposure risk. However, no study to date has characterized how these viruses evolve during such outbreaks in nature, leaving key gaps in our understanding of viral evolution at the human animal interface. In this study, we generated replicate, within-sample diversity data from naturally infected domestic birds, and used these data to quantify viral diversity and local spread. We show that Pennsylvania outbreaks were supported by repeated introductions into the state, sparking multiple co-circulating transmission chains. Live bird market outbreaks in the same place and time were seeded independently, suggesting that these markets experience risk from varied sources, and may benefit from enhanced biosecurity and tracking. These markets also support high circulating diversity, viral mixing between species, and outbreaks that cross state lines, highlighting the need to better control transmission in these settings. We find that within-sample diversity was frequently shared among samples from the same outbreak and can be used to clarify patterns of host-specific variant sharing in outbreak settings. These data imply high circulating diversity within these outbreaks that could potentially be used to enhance transmission linkage, and to detail differences in outbreak flow between commercial and LBMs. Finally, while rare, we find that a small subset of known adaptive mutations are present in samples from domestic bird outbreaks. These data confirm that putative adaptive variants may be present in settings where humans and animals interact, providing evolutionary opportunities for viral adaptation and spillover.

Our data identified repeated independent introductions into Pennsylvania, consistent with pervasive viral flow among US states. We present compelling evidence of transmission between live bird market settings and distributors in New York, Pennsylvania, and New Jersey, suggesting that viral flow within these systems may be occurring across state lines. The independent introductions of LBM outbreaks in the same county 3 weeks apart further suggest that these markets are at risk from varied sources, including potential spillover from wild birds, transmission of sick birds among markets, and infections from bird transport and holding centers. The diverse geographic origins of birds that flow through these markets, the propensity to mix species together, and historical lack of regulations and compliance to biosecurity measures collectively make live bird markets high risk for influenza transmission ^59,60^. Since the influx of HPAI in North America, stronger regulations and biosecurity measures have been implemented at the state level for live bird market systems, in particular by the Live Bird Marketing System Biosecurity Assurance Program ^61^. Our findings, along with ongoing detections in the LBMs and the presence of putative adaptive mutations, suggest that additional interventions, improvements, or regulation may be needed.

We find no evidence in our study that the outbreaks we analyzed were connected to each other. This finding is particularly relevant for the case of the LBM outbreaks, which occurred in the same location less than a month apart. While our dataset involves only a small fraction of all domestic outbreaks in Pennsylvania, these results are encouraging, and suggest that to some degree, biosecurity and outbreak containment strategies are successfully reducing farm to farm spread. Still, our data show that viral diversification does occur during these outbreaks, and that in some cases, known adaptive variants are present. Within-sample data further confirmed that variants are frequently shared among distinct host species within LBMs, implying a high degree of viral mixing between species in these settings. While this diversity and variants arising in these infections seem to be primarily confined to these outbreaks, it is concerning that viral diversity is being propagated and subsequently transmitted within farms, particularly because these interfaces are those at highest risk for human exposure. Given the finding of PB2 D701N and the high rate of variant sharing we find in these settings, these results suggest a heightened risk of domestic outbreaks as a source of adaptive variation. Our data support continuing to stamp out transmission chains to reduce the likelihood of viral adaptation.

We find some phylogenetic evidence that Anseriformes (ducks and geese) are basal to Galliformes (chickens, chukars, guineas) in two commercial outbreak clusters (February 2023, B3.3 and July 2025, D1.1). While our sample size is very small, these data align with numerous reports from Asia of ducks becoming infected first and subsequently infecting chickens in domestic settings ^62,63^. Given that waterfowl are the primary reservoir of avian influenza viruses and thus are more frequently asymptomatic, this “trojan horse” effect could apply to outbreaks in North America as well ^63,64^. Future studies that focus more intensely on verifying whether this pattern occurs across outbreaks more broadly could be useful for designing biosecurity protocols for mixed species farms, or duck and chicken farms in close proximity, which may require stricter biosecurity to limit interaction between these host groups.

To our knowledge, our study provides the first data on within-sample diversity in natural infections of domestic birds in North America. We found that diversity is generally low within-sample, with comparable numbers of iSNVs compared to previous studies of influenza in natural human infections, which report fewer than 10 iSNVs/sample on average ^17,57,65,66^. However, we do find a higher frequency of mid to high frequency variants, and a shift in variant frequencies away from very low frequency variants (<10%). These differences likely arise from the pooling in our samples, which may combine within-host and consensus variants among infections together. Because of sample pooling, iSNV counts in our dataset could either overestimate (by grouping together multiple unique infections and their variants), or could underestimate (by diluting out minority variants below our iSNV calling thresholds) iSNV numbers. Given that we find no difference in iSNV between Anseriformes and Galliformes, which are pooled differently, it is also possible that these pooling protocols have little impact on diversity metrics. However, we do find that D1.1 samples harbored more within-sample variants, which could arise from D1.1’s rapid spread and propensity to infect more diverse hosts, or from differences in infection dynamics ^43,67^. The degree to which the higher diversity we measure is due to inherent differences in viral replication dynamics, shedding dynamics, or epidemiologic factors like the transmission rate or serial interval, is not currently clear.

Regardless of why, our data suggest that samples from the same outbreak share substantial within-sample variation, providing a genetic signature of epidemiologic linkage. Previous studies of RNA virus transmission in both household cohorts and laboratory settings have found influenza transmission bottlenecks to be very narrow, limiting the levels of variants being shared between individuals ^17,18,20,68,69^. Prior work that has attempted to use within-host variation to study human infections among known transmission pairs have found success in settings with high diversity and loose bottlenecks (e.g., HIV, HCV) ^11,13,70^, but have not worked for respiratory viruses due to the tight transmission bottleneck. While we cannot measure bottleneck size with our data due to sample pooling, our data suggest that samples collected from these agricultural settings retain much higher frequencies of shared variants than expected. This pattern could arise if transmission bottlenecks in these agricultural settings are much looser than previously seen in human or animal transmission pairs, which could be explained by high-density indoor poultry rearing, a high force of infection, or infection via environmental or water contamination that enables a high inoculum ^71–73^. Alternatively, the sample pooling procedure itself could result in substantial shared variation simply by combining co-circulating variants together. Regardless, our data suggest that within-sample variation of pooled, domestic samples retains useful information about transmission linkage that could be used to augment transmission inference. As these viruses continue to circulate in North American wild birds and spill into agricultural settings, resolving the source of these outbreaks could be useful for designing functional and effective outbreak prevention strategies.

Our study has important limitations. Sequences from wild animal infections beyond February 2025 are limited, which poses a central challenge for local-scale outbreak tracking. Given the high frequency of outbreaks that arise from spillover, wild bird data is increasingly critical for disentangling sources of HPAI outbreaks and to rule out transmission between agricultural and production settings ^3^. An additional limitation of our dataset is the pooling of outbreak samples, which are variable and sometimes unclear. While sample pooling likely increases testing efficiency, it precludes direct identification of variant frequency and source within individual animals, making inference of parameters like bottleneck size impossible.

Future, targeted studies that directly aim to address this gap may improve inference of on-farm transmission parameters. Finally, we only characterized a small subset of outbreaks in Pennsylvania, which varied across production types ^74–76^. Farm to farm variation in the level of regulation, biocontrol, and stringent husbandry practices is likely a key indicator for outbreak risk that is impossible to study currently due to a lack of metadata. These factors of species, production type, and density could certainly impact transmission and should be accounted for in models and future analyses when possible ^77^.

Taken together, this study produced first of its kind data from natural infections in domestic birds. By applying a series of phylogenetic and within-sample approaches, we quantify diversity, evolution, and transmission across domestic outbreaks. The multiple independent introductions into the state throughout the epizootic show that Pennsylvania is likely a useful model state for understanding finer-scale transmission dynamics of H5N1. Our results present evidence that high levels of diversity is being shared within domestic outbreaks, and particularly that continued surveillance and increased regulation of live bird market systems is important for controlling interstate spread particularly within the Northeastern United States. Future study and surveillance where samples are collected from individual domestic infections will allow for true quantification of transmission bottlenecks to further inform biosecurity and control measures in domestic settings.

## Acknowledgments

This work was supported by the Penn Center for Research on Coronaviruses and Other Emerging Pathogens, the Pennsylvania Department of Agriculture, and by the National Institute of Allergy and Infectious Diseases, National Institutes of Health, Department of Health and Human Services (co. number 75N93021C00015) and by grant number R01AI195533. L.H.M. is a Pew Biomedical Scholar and is supported by funding from the Pew Charitable Trusts.

## Materials & Methods

### Ethics Statement

This study used residual diagnostic samples/specimens that were originally submitted to the Pennsylvania Animal Diagnostic Laboratory System for routine diagnostic testing. No animals were handled or sampled specifically for this study.

### Diagnostic testing and RNA extraction

Diagnostic testing was performed at the Pennsylvania Animal Diagnostic Laboratory System (PADLS), a National Animal Health Laboratory Network (NAHLN)-approved laboratory. Influenza A virus detection and subtype identification were performed using NAHLN -approved real-time reverse transcription PCR assays according to National Veterinary Services Laboratory (NVSL; Ames, IA, USA)-approved protocols. Briefly, nucleic acid was extracted using a commercial kit (MagMax-96 Viral RNA Isolation kit; Thermo Fisher) according to the manufacturer’s instructions. Initially, the samples were screened for IAV by rRT-PCR targeting the matrix gene. All matrix-non-negative samples were subsequently tested using H5 subtype assays, including assays targeting H5 viruses associated with clade H5 2.3.4.4b. Additionally, all IAV-non-negative samples were sent to NVSL for confirmatory testing and further characterization.

### Sample pooling

Samples were originally collected following the pooling guidelines outlined for pathogen detection by the National Veterinary Services Laboratories (NVSL) ^28^. Specific pool numbers and swab type for each sample can be found in Table 1. Given that these are pooled individual swabs, we cannot be certain or quantify how many individual positives are represented in each pool. For example, a chicken sample of 10 swabs could represent 10 infected individuals with equal viral concentrations or could be 2 infections with varying concentrations and multiple negative swabs. For these reasons, we describe sub-consensus variants (iSNVs) as within sample variants, and interpret them as proxies for the amount of circulating diversity (both within and between host) present in the sample pool.

### cDNA generation and PCR amplification

cDNA was generated from viral RNA (vRNA) using the Protoscript II first strand synthesis kit (NEB, catalogue # E6560L) and the Uni12 primer (AGCRAAAGCAGG). When sufficient vRNA volume permitted (>8 μl), two separate reactions were conducted per vRNA sample and were treated as complete technical replicates to control for downstream potential PCR and sequencing errors ^29^. The complete reverse transcription protocol can be found here: https://github.com/moncla-lab/h5-sequencing-protocol-dev/blob/main/RT_and_PCR_protocol.md. Single stranded complementary DNA was then used as a template for PCR amplification of all eight genes. Primers were adapted from H1N1 primers from ^20^. These primers utilize the Hoffman universal primer sequences ^78^ with several bases added for specificity based on alignments of relevant H5N1 sequences to improve specificity. Additional internal primers have been added for polymerase genes and HA to improve sequencing coverage (see table below). PCR was performed with the Q5 Hotstart DNA polymerase (NEB, catalogue # M0493L). Following PCR amplification, individual amplicons were cleaned via a 1X bead cleanup using AMPure XP beads and 1µl of each sample was used to quantify using Qubit 1X dsDNA HS Assay kit (Thermo Fisher, catalogue #Q33230). PCR for technical replicates was performed in separate experimental batches. Negative controls (water) were included for each reverse transcription experiment, and each separate PCR experiment and were sequenced along with samples.

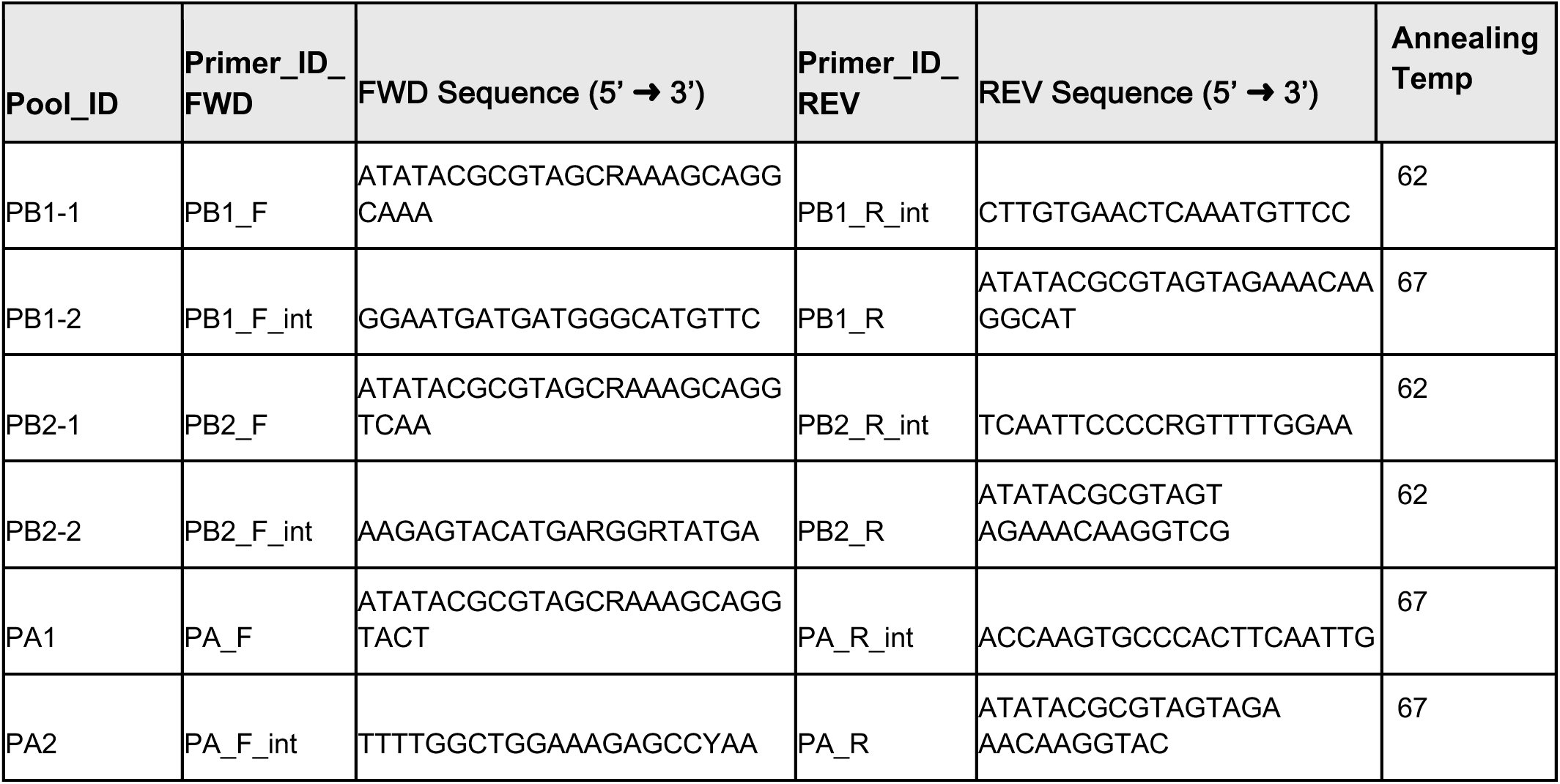
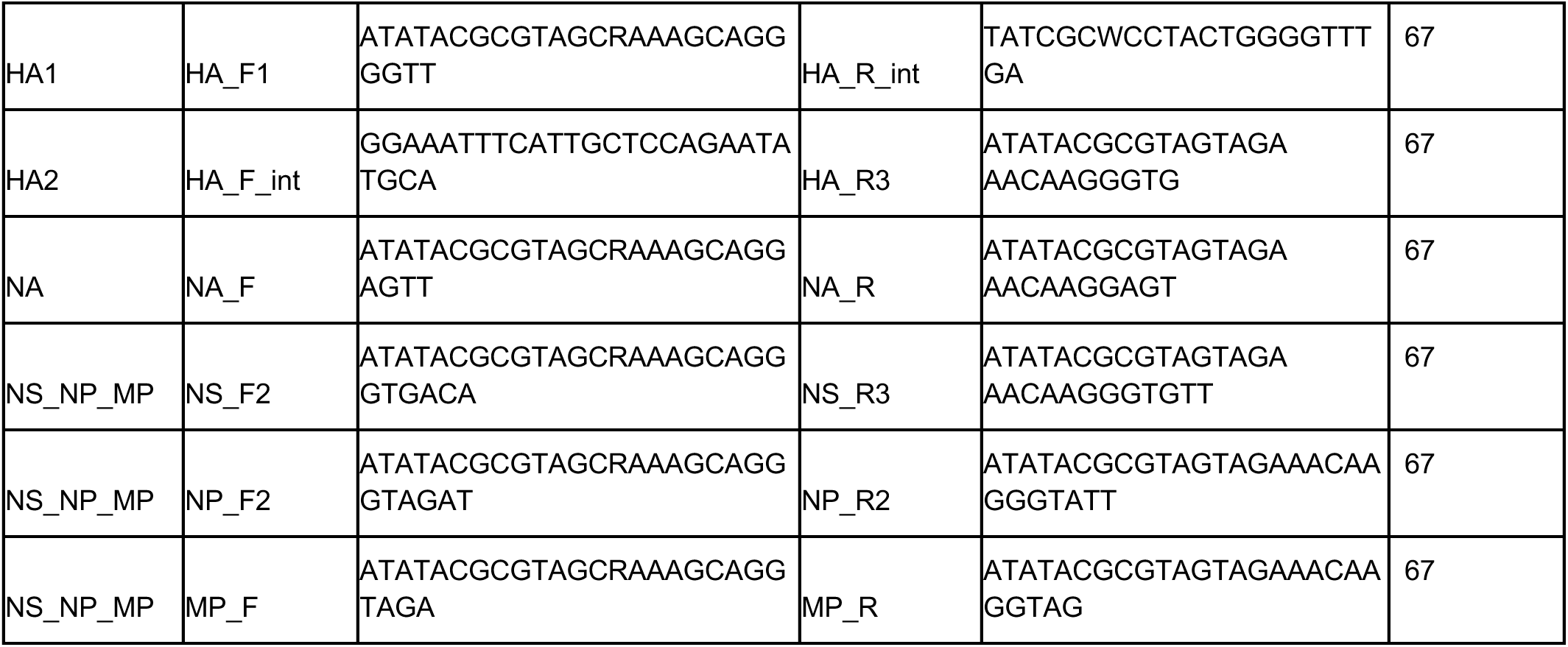

### Library Preparation and Sequencing

PCR amplicons were diluted, requantified by Qubit, and all amplicons originating from the same samples were pooled in equimolar ratios to a final concentration of 0.2 ng/5 µl. Library preparation was then performed with the Nextera XT DNA Library Prep kit (Illumina, catalogue # FC-131-1096), following the manufacturer’s recommended protocol. Specifically, each sample or genome was enzymatically fragmented and tagged with short oligonucleotide adapters, followed by fifteen cycles of PCR for template indexing. Libraries were then purified by two consecutive Ampure bead cleanups ((0.5X and 0.7X) and were quantified by Qubit. The average sample fragment length and purity were then determined on the Agilent Tapestation using the D1000 or D5000 High Sensitivity ScreenTape Analysis. The complete library preparation protocol can be found here: https://github.com/moncla-lab/h5-sequencing-protocol-dev/blob/main/Library_preparation.md

Prepared libraries were pooled in equimolar concentrations to a final concentration of 4nM, and run on a MiniSeq Miniseq High or Mid output kit for 300 cycles (2 × 150 bp paired-end) (Illumina, catalogue # FC-420-1003, or #FC-420-1004). Demultiplexed files were output in FASTQ format. Samples were sequenced across six independent sequencing runs.

### Raw Sequence Data Processing and Variant Calling

Raw FASTQ files were processed using custom bioinformatic pipelines, software and documentation available on GitHub here: https://github.com/moncla-lab/illumina-pipeline. Briefly, all sequencing experiments for a given sample were merged into a single forward and reverse FASTQ files based on the input metadata sample manifest. Paired-end reads of a minimum read length of 50, and an average read quality score of Q30 within a 5 base pair sliding trimming window were trimmed for each replicate using Trimmomatic ^79^. Paired-end reads were then merged and mapped to a custom H5N1 reference sequence with appended full UTRs using BowTie2 ^80^. Consensus sequences for each replicate were called by reporting the majority base at each site with a base calling cut-off of 25X coverage. For each sample, replicate consensus were merged to report out a sample-level consensus and any discrepancies were reported. Minority variants were called against their replicate consensus sequence independently for technical replicates using Samtools ^81^ and Varscan ^82^ using a variant frequency threshold of 2%, minimum 100X coverage.

### Negative controls

Negative controls (H20) were included in each RT and PCR experiment, and were processed as if they were true samples. Each negative control was sequenced along with samples. For negative controls, we confirmed lack of laboratory contamination if read depth was under 10X. In one sequencing run, we had negative controls with reads above 10X coverage in one segment (PA). To account for this potential contamination, exact regions with high read depth in negative controls were identified using the depth.txt output from the illumina-pipeline, and exact samples associated with preparation were confirmed from our experiment notes (see example sheets in https://github.com/moncla-lab/hpai-pa/lab-experiments). The amplicon with high reads spanned PA 999-2233 and was run with samples c_com1, c_com2, and c_com3. All variants called in this region of PA from these samples processed with negative control with potential contamination were removed from analysis. This included 8 single-replicate variants from these three samples.

### Variant and Within-sample Diversity Analysis

As described above, within-sample variants were called at a minimum coverage of 100X, and a minimum of 2% frequency. Only variants called in both replicates were used for analyses. Variant frequency was averaged across replicates. All code for variant analysis is available at https://github.com/moncla-lab/hpai-comm-pa.

#### Depth normalization

To normalize sequencing coverage in analysis of SNVs per sample, as well as for the shared variant site permutation test, we normalized variant coverage by sequencing depth. Here, we utilized the output from our analysis pipeline’s depth.txt files for each sample and each replicate which outputs the depth of coverage at each base. We merged these depth reports for each replicate and quantified the number of bases for each gene where sequence coverage was >100X for both replicates. The proportion coverage for each gene was then used to normalize the variant counts for each sample for each gene by the number of nucleotides of coverage sufficient to call variants, normalized per 1000 bases. For example, if a sample had 9,000 bases with >100X coverage and 9 variants called for the sample, the depth normalized variant count for the sample would be 1 variant/1kb of coverage.

#### Shared sites permutation analysis

To better determine whether the observed level of sharing was higher than expected by chance, we adapted the approach described in Moncla et al ^35^ to conduct a permutation test. We simulated assigning the same number of mutations that we had in our actual data to the same number of samples, but randomly assigning the position of these mutations. Depth of sequencing coverage was accounted for for each sample to restrict placement of variants, such that variants could only be placed at sites with at least 100x coverage. Simulations were run in 20,000 iterations for each individual cluster for which there were samples with within-host variants (1-6), as defined by date and location. For each simulation, the total number of shared variant sites was tabulated and the distribution across the 20,000 iterations was plotted for each cluster against the observed number of shared sites. This permutation test was also repeated to estimate the expected number of shared variant sites between clusters in 100,000 iterations.

For this second analysis, the number of variants and sequencing coverage per gene per sample were once again taken into account. For each simulation, this true number of mutations were assigned for each sample but at random positions. These mutations were then pooled by cluster and compared to the randomly assigned mutations for each sample in other clusters. The number of mutations at shared sites between clusters were then tabulated for each simulation and the distribution of these shared variants counts between clusters were plotted for the 100,000 iterations against the observed number of shared sites from the data.

#### Linear regression models

To determine the relative contributions of domestic status, host order, genotype, and Ct-value to predicting the number of variants per sample, we fit a linear regression model to the data in python using the statsmodel package ^83^. We modelled the depth normalized iSNVs per sample per 1kb coverage as our outcome variable, and evaluated the contribution of four predictor variables on this outcome: domestic status, host order, genotype, and Ct. For binary traits (domestic status, host order, and genotype), we coded the values as follows: Galliforme = 1, Anseriforme = 0; commercial = 1, LBM = 0; D1.1 = 1, B3.3 = 0. Ct-values were normalized by subtracting the mean Ct-value, divided by the standard deviation, from each value to reduce multicollinearity in the model. The full model was defined as follows: normalized SNVS per sample per 1 kb coverage ∼ host order galliformes + domestic status commercial + genotype D1.1 + normalized Ct-value.

To determine the contributing factors for the proportion of shared variants between samples, we similarly fit a linear regression model. Here, for our outcome variable, we performed pairwise comparisons for each pair of samples to compute their proportion of shared variants with each other sample. For example, for samples A and B, we calculated the proportion of SNVs in sample B that were present in sample A, as well as the proportion of SNVs in sample B that were in sample A. We calculated these proportions in both directions for all sample combinations. We included four binomial predictor variables: whether the pair was from the same cluster, same host, same genotype, and same domestic status with 1 representing the same and 0 representing different for each variable. The model was as follows: proportion of shared variants ∼ same cluster + same host order + same domestic status.

#### Functional sites

To identify within-sample variants at previously annotated functional sites, we compiled a database of annotated sites from multiple sources ^35–37^, confirmed H5 numbering, and cross-referenced these positions against our data. For HA mutations, these were restricted to H5N1 mutations only. For NA mutations, these sites were restricted to N1 mutations only. All internal genes considered all IAV subtypes. Nonsynonymous variants at these sites were then manually investigated to confirm whether the exact amino acid change previously annotated was present.

### Phylogenetic reconstruction and analysis

We downloaded all currently available H5Nx virus genomes from the EpiFlu Database of the Global Initiative for Sharing All Influenza Data (GISAID, https://www.gisaid.org/) and added consensus genomes from all of our sequenced Pennsylvania samples. Metadata was cleaned and organized to include, when possible, migratory flyway, species and order, as well as domestic status. Sequences and metadata were then processed using Nextstran’s augur software ^84^. Sequences were filtered by length to remove short sequences for each segment. We additionally excluded all sequences prior to 2021; sequences for which the host was unknown, laboratory derived, environmental, or ferret; sequences not from North America or unknown country or region; and all clades other than 2.3.4.4b as designated by LABEL ^85^. We included all sequences from Pennsylvania, and otherwise subsampled 50 sequences per group based on month, host, and location. Sequences for each gene were aligned using MAFFT ^86^.

We used the A/goose/Czech Republic/18520-2/2021(H5N1) genome to define reference coordinates (Genbank accession numbers: OL638145-OL638152). Using the Nextstrain workflow, a maximum likelihood phylogeny was inferred using IQ-TREE ^87^, and the molecular clock was inferred with TreeTime ^88^. TreeTime was next used to infer the ancestral trait reconstruction at internal nodes. We inferred these internal node traits for the trait “border_states” with three defined states of “Pennsylvania”, “border”, or “non-PA”. Border states are: New York, New Jersey, Delaware, Maryland, West Virginia, and Ohio. Our final trees are available here (https://github.com/moncla-lab/hpai-comm-pa/tree/main/phylo), and include the following number of sequences for the HA tree: 6,666 total sequences (243 Pennsylvania, 452 border, 5,971 non-border). Sequence counts for other segments were comparable.

As a proxy for quantifying introductions and exportations to and from Pennsylvania, we calculated the transitions between states across branches of phylogenies from ancestral trait reconstructions on the trait “border state” using the Baltic python package (https://github.com/evogytis/baltic). Introductions were quantified using all eight gene trees and presented in results as an average across all eight gene trees. The resulting HA tree is visualized in the main text, and additional gene trees are available in Supplementary (Figure 2).

To further investigate transmission cluster dynamics, we constructed concatenated full genome trees for B3.3 and D1.1 genotype sequences. Genotypes were first assigned to all sequences in our North American dataset using the USDA Genoflu tool ^1^(https://github.com/moncla-lab/GenoFLU-multi). Gene sequences that were assigned B3.3 and D1.1 genotypes were then concatenated into full genome sequences and a genotype-specific full-genome maximum-likelihood trees were then run using the previously described Nextstrain pipeline containing 60 B3.3 full genome sequences and 661 D1.1 full genome sequences. For visualization, divergence trees in Newick format were exported from Nextstrain and plotted using Baltic (https://github.com/evogytis/baltic).

#### Bayesian phylogenetic analysis of D1.1 full genome sequences

To infer rates of transition between domestic Pennsylvania and New York, we utilized a Bayesian phylogenetic reconstruction in BEAST (v1.10.4) ^89^. Here we utilized a 315 concatenated full genome sequence alignment selected from the maximum likelihood phylogenetic tree of full D1.1 genomes which formed a distinct cluster including the two Pennsylvania live bird market outbreaks. We constructed a trait called “pa_ny_domstat” which classified strains as Pennsylvania domestic (PA_domestic, which includes LBM), New York domestic (NY_domestic), all other domestic sequences from other states (domestic), all wild sequences from any state (wild), and removed any sequences with unknown domestic status or unknown dates from the alignment which resulted in a 215 sequence alignment (51 PA domestic; 45 NY domestic; 35 domestic; 84 wild sequences). For our discrete trait model, we implemented an asymmetric discrete trait model and implemented the Bayesian stochastic search variable selection (BSSVS) to infer transition rates and support of these estimates ^90^.

Bayes Factors were calculated for quantifying support of these transition rates using Spread3 v.0.9.6 ^91^. In addition to quantifying discrete trait rates, we enumerated the total number of jumps across the posterior using Markov Jumps and Rewards, computing jumps for transitions including PA domestic and NY domestic (omitting transitions between other wild and domestic), and rewards for measuring the persistence time for these discrete trait states (time in PA domestic and time in NY domestic) in the phylogeny ^92,93^.

We used the following priors and settings. We used an HKY nucleotide substitution model with gamma distributed rate variation among nucleotide sites. We used a log-normal relaxed molecular clock model with a prior mean value of .00515 based on the molecular clock rate in our maximum likelihood D1.1 full genome phylogeny. We ran four independent MCMC chains with a chain length of 100 million states logging every 100,000 states. We checked results in Tracer v1.7.2 to ensure sufficient effective sample size (ESS) of over 200 and reasonable estimates for all parameters ^94^. We combined tree files from each independent iteration removing 10% burn-in to get a tree file with 1000 posterior trees using Logcombiner v.1.10.4 ^89^. A maximum clade credibility (MCC) tree was then constructed using TreeAnnotator v.1.10.4 and used for visualizations and for summarizing the posterior probabilities for inferred traits ^89^. XML files were constructed first in BEAUTI and then manually edited to include Markov jumps and rewards. XML, and output files are available here: https://github.com/moncla-lab/hpai-comm-pa/tree/main/beast.

## Data Availability

All raw sequence data are available in the SRA under Bioproject PRJNA1484614. All other data that were used in this analysis were sourced from public databases (NCBI) and a list of these accession numbers are available in our github repository https://github.com/moncla-lab/hpai-comm-pa/blob/main/ncbi_sequences.tsv. The acknowledgement table for GISAID isolates used in this analysis is provided in Supplementary Table 2, and also available in the github repository https://github.com/moncla-lab/hpai-comm-pa. These sequences are additionally available in the EPI_SET_260701sb, with an access link provided in Supplementary Appendix 1. The complete reverse transcription, PCR protocol, including primer sequences, pooling scheme, and annealing temperatures, library preparation, and example experiment spreadsheets can be found here: https://github.com/moncla-lab/h5-sequencing-protocol-dev. The deep-sequencing analysis pipeline used for this study is available at https://github.com/moncla-lab/illumina-pipeline. All code used to analyze and plot these data, as well as associate data files for within-host variant calls and phylogenetic trees are available at https://github.com/moncla-lab/hpai-comm-pa.

## Referencesm

1. Youk, S. et al. H5N1 highly pathogenic avian influenza clade 2.3.4.4b in wild and domestic birds: Introductions into the United States and reassortments, December 2021–April 2022. Virology 587, 109860 (2023).

2. Caliendo, V. et al. Transatlantic spread of highly pathogenic avian influenza H5N1 by wild birds from Europe to North America in 2021. Sci. Rep. 12, 11729 (2022).

3. Damodaran, L., Jaeger, A. S. & Moncla, L. H. Ecology and spread of the North American H5N1 epizootic. Nature 649, 432–441 (2026).

4. Klaassen, M. & Wille, M. The plight and role of wild birds in the current bird flu panzootic. *Nat*. Ecol. Evol. 7, 1541–1542 (2023).

5. Lee, D.-H. et al. Transmission Dynamics of Highly Pathogenic Avian Influenza Virus A(H5Nx) Clade 2.3.4.4, North America, 2014–2015. Emerg. Infect. Dis. 24, 1840–1848 (2018).

6. Hicks, J. T. et al. Agricultural and geographic factors shaped the North American 2015 highly pathogenic avian influenza H5N2 outbreak. PLoS Pathog. 16, e1007857 (2020).

7. Dholakia, V. et al. Polymerase mutations underlie early adaptation of H5N1 influenza virus to dairy cattle and other mammals. Nat. Commun. 17, 1603 (2026).

8. Panova, A. S. et al. Genotype A3 influenza A(H5N1) isolated from fur seals shows high virulence in mammals, but not airborne transmission. Sci. Rep. 15, 44463 (2025).

9. Dadonaite, B. et al. Deep mutational scanning of H5 hemagglutinin to inform influenza virus surveillance. PLoS Biol. 22, e3002916 (2024).

10. Signore, A. V. et al. Spatiotemporal reconstruction of the North American A(H5N1) outbreak reveals successive lineage replacements by descendant reassortants. Sci. Adv. 11, eadu4909 (2025).

11. Thompson, R. N. et al. Link between the numbers of particles and variants founding new HIV-1 infections depends on the timing of transmission. Virus evolution 5, vey038 (2019).

12. Wang, G. P., Sherrill-Mix, S. A., Chang, K.-M., Quince, C. & Bushman, F. D. Hepatitis C virus transmission bottlenecks analyzed by deep sequencing. J. Virol. 84, 6218–6228 (2010).

13. Li, H. et al. Single-genome sequencing of hepatitis C virus in donor-recipient pairs distinguishes modes and models of virus transmission and early diversification. J. Virol. 90, 152–166 (2016).

14. Haaland, R. E. et al. Inflammatory genital infections mitigate a severe genetic bottleneck in heterosexual transmission of subtype A and C HIV-1. PLoS pathogens 5, e1000274 (2009).

15. Holmes, K. E. et al. Viral expansion after transfer is a primary driver of influenza A virus transmission bottlenecks. PLoS Biol. 23, e3003352 (2025).

16. Varble, A. et al. Influenza A virus transmission bottlenecks are defined by infection route and recipient host. Cell Host Microbe 16, 691–700 (2014).

17. McCrone, J. T. et al. Stochastic processes constrain the within and between host evolution of influenza virus. eLife 7, (2018).

18. Braun, K. M. et al. Acute SARS-CoV-2 infections harbor limited within-host diversity and transmit via tight transmission bottlenecks. PLoS Pathog. 17, e1009849 (2021).

19. Sinclair, P., Zhao, L., Beggs, C. B. & Illingworth, C. J. R. The airborne transmission of viruses causes tight transmission bottlenecks. Nat. Commun. 15, 3540 (2024).

20. Braun, K. M. et al. Avian H7N9 influenza viruses are evolutionarily constrained by stochastic processes during replication and transmission in mammals. Virus Evol. 9, vead004 (2023).

21. Leyson, C. M., Criado, M. F., Youk, S. & Pantin-Jackwood, M. J. Low pathogenicity H7N3 avian influenza viruses have higher within-host genetic diversity than a closely related high pathogenicity H7N3 virus in infected turkeys and chickens. Viruses 14, 554 (2022).

22. Iqbal, M. et al. Within-host variation of avian influenza viruses. Philos. Trans. R. Soc. Lond. B Biol. Sci. 364, 2739–2747 (2009).

23. CDC. USDA reported H5N1 bird flu detections in poultry. Avian Influenza (Bird Flu) https://www.cdc.gov/bird-flu/situation-summary/data-map-commercial.html (2025).

24. USDA - National Agricultural Statistics Service - Charts and Maps - Layers and Eggs: Production by State, US. https://www.nass.usda.gov/Charts_and_Maps/Poultry/eggmap.php.

25. Confirmations of Highly Pathogenic Avian Influenza in Commercial and Backyard Flocks. Animal and Plant Health Inspection Service https://www.aphis.usda.gov/livestock-poultry-disease/avian/avian-influenza/hpai-detections/commercial-backyard-flocks (2023).

26. Jagne Dvm Dacpv, J. F., Bennett Dvm Mph Dacvpm, J. & Collins Dvm, E. Live bird markets of the Northeastern United States. Dela. J. Public Health 7, 52–56 (2021).

27. Pennsylvania Bulletin. https://www.pacodeandbulletin.gov/Display/pabull?file=/secure/pabulletin/data/vol56/56-6/193.html&search=1&searchunitkeywords=live,bird,market.

28. Torchetti, M. Sample Collection for Avian Influenza and Newcastle Disease. https://www.aphis.usda.gov/sites/default/files/avian-sample-collection-ai-newcastle.pdf (2024).

29. McCrone, J. T. & Lauring, A. S. Measurements of intrahost viral diversity are extremely sensitive to systematic errors in variant calling. J. Virol. 90, 6884–6895 (2016).

30. Xue, K. S. & Bloom, J. D. Reconciling disparate estimates of viral genetic diversity during human influenza infections. Nat. Genet. 51, 1298–1301 (2019).

31. Tewari, D. et al. Genotype diversity of highly pathogenic avian influenza H5N1 clade 2.3.4.4b in Pennsylvania poultry during disease outbreak from April 2022 to march 2023. Viruses 18, 502 (2026).

32. Dudas, G. & Bedford, T. The ability of single genes vs full genomes to resolve time and space in outbreak analysis. BMC Evol. Biol. 19, 232 (2019).

33. Imai, M. et al. Experimental adaptation of an influenza H5 HA confers respiratory droplet transmission to a reassortant H5 HA/H1N1 virus in ferrets. Nature 486, 420–428 (2012).

34. Linster, M. et al. Identification, characterization, and natural selection of mutations driving airborne transmission of A/H5N1 virus. Cell 157, 329–339 (2014).

35. Moncla, L. H. et al. Quantifying within-host diversity of H5N1 influenza viruses in humans and poultry in Cambodia. PLoS Pathog. 16, e1008191 (2020).

36. Nguyen, T.-Q. et al. Emergence and interstate spread of highly pathogenic avian influenza A(H5N1) in dairy cattle in the United States. Science 388, eadq0900 (2025).

37. Mojsiejczuk, L., et al. Flu Mutation Explorer: An interactive platform for mapping host adaptation mutations in influenza A viruses. bioRxiv 2026.07.22.740012 (2026) doi:10.64898/2026.07.22.740012.

38. Banyard, A. C. et al. Detection of clade 2.3.4.4b H5N1 high pathogenicity avian influenza virus in a sheep in Great Britain, 2025. Emerg. Microbes Infect. 14, 2547730 (2025).

39. Bruno, A., de Mora, D., Garcia-Bereguiain, M. A. & Cristina, J. Phylogenetic and mutation analysis of hemagglutinin gene from Highly Pathogenic Avian Influenza Virus H5 clade 2.3.4.4b in South America. Viruses 17, 924 (2025).

40. Fan, S. et al. Novel residues in avian influenza virus PB2 protein affect virulence in mammalian hosts. Nat. Commun. 5, 5021 (2014).

41. Steel, J., Lowen, A. C., Mubareka, S. & Palese, P. Transmission of influenza virus in a mammalian host is increased by PB2 amino acids 627K or 627E/701N. PLoS Pathog. 5, e1000252 (2009).

42. Czudai-Matwich, V., Otte, A., Matrosovich, M., Gabriel, G. & Klenk, H.-D. PB2 mutations D701N and S714R promote adaptation of an influenza H5N1 virus to a mammalian host. J. Virol. 88, 8735–8742 (2014).

43. Crespo-Bellido, A. et al. Emergence of D1.1 reassortant H5N1 avian influenza viruses in North America. bioRxiv 2025.12.19.695329 (2025) doi:10.64898/2025.12.19.695329.

44. Quirk, G. E. et al. Variable transmission efficiency of mammalian origin HPAI D1.1 H5N1 strains in ferrets. bioRxivorg 2026.05.07.722809 (2026) doi:10.64898/2026.05.07.722809.

45. Sun, B. et al. Emergence of the novel PA-D27G mutation conferring reduced baloxavir susceptibility in influenza A viruses circulating in China, 2018-2025. Emerg. Microbes Infect. 15, 2620222 (2026).

46. Ping, J. et al. Genomic and protein structural maps of adaptive evolution of human influenza A virus to increased virulence in the mouse. PLoS One 6, e21740 (2011).

47. Yang, J. et al. The haemagglutinin gene of bovine-origin H5N1 influenza viruses currently retains receptor-binding and pH-fusion characteristics of avian host phenotype. Emerg. Microbes Infect. 14, 2451052 (2025).

48. Kim, W.-J., Hur, K.-Y., Park, H. W., Lee, S.-W. & Yoo, J.-Y. Pathogenicity of the H1N1 influenza virus enhanced by functional synergy between the NPV100I and NAD248N pair. PLoS One 14, e0217691 (2019).

49. Chothe, S. K. et al. Marked neurotropism and potential adaptation of H5N1 clade 2.3.4.4.b virus in naturally infected domestic cats. Emerg. Microbes Infect. 14, 2440498 (2025).

50. Chen, C. et al. Highly pathogenic avian influenza A H5N1 virus infection in an immunocompromised domestic cat. ASM Case Rep. 1, e00134–25 (2025).

51. Boltz, D. A. et al. Emergence of H5N1 avian influenza viruses with reduced sensitivity to neuraminidase inhibitors and novel reassortants in Lao People’s Democratic Republic. J. Gen. Virol. 91, 949–959 (2010).

52. Yao, Q. et al. Emergence and evolution of novel canine-avian reassortant H3N2 influenza A viruses in duck in Leizhou Peninsula, China. Front. Microbiol. 13, 857800 (2022).

53. Adams, S. E. et al. Effect of influenza H1N1 neuraminidase V116A and I117V mutations on NA activity and sensitivity to NA inhibitors. Antiviral Res. 169, 104539 (2019).

54. Zhang, J. et al. The D253N mutation in the polymerase basic 2 gene in avian influenza (H9N2) virus contributes to the pathogenesis of the virus in mammalian hosts. Virol. Sin. 33, 531–537 (2018).

55. Mok, C. K. P. et al. Amino acid residues 253 and 591 of the PB2 protein of avian influenza virus A H9N2 contribute to mammalian pathogenesis. J. Virol. 85, 9641–9645 (2011).

56. Wang, X. et al. Amino acid mutations PB1-V719M and PA-N444D combined with PB2-627K contribute to the pathogenicity of H7N9 in mice. Vet. Res. 55, 86 (2024).

57. Debbink, K. et al. Vaccination has minimal impact on the intrahost diversity of H3N2 influenza viruses. PLoS Pathog. 13, e1006194 (2017).

58. Bendall, E. E. et al. Influenza A virus within-host evolution and positive selection in a densely sampled household cohort over three seasons. Virus Evol. 10, veae084 (2024).

59. Cardona, C., Yee, K. & Carpenter, T. Are live bird markets reservoirs of avian influenza? Poult. Sci. 88, 856–859 (2009).

60. Bulaga, L. L. et al. Descriptive and surveillance studies of suppliers to New York and New Jersey retail live-bird markets. Avian Dis. 47, 1169–1176 (2003).

61. Pennsylvania Bulletin. https://www.pacodeandbulletin.gov/Display/pabull?file=/secure/pabulletin/data/vol55/55-39/1331.html.

62. Kwon, J.-H. et al. Domestic ducks play a major role in the maintenance and spread of H5N8 highly pathogenic avian influenza viruses in South Korea. Transbound. Emerg. Dis. 67, 844–851 (2020).

63. Hulse-Post, D. J. et al. Role of domestic ducks in the propagation and biological evolution of highly pathogenic H5N1 influenza viruses in Asia. Proc. Natl. Acad. Sci. U. S. A. 102, 10682–10687 (2005).

64. Wu, S. X., Davis, C. N., Arnold, M. & Tildesley, M. J. The role of ducks in detecting Highly Pathogenic Avian Influenza in small-scale backyard poultry farms. PLoS Comput. Biol. 22, e1013357 (2026).

65. Sobel Leonard, A., et al. Within-host influenza viral diversity in the pediatric population as a function of age, vaccine, and health status. Virus Evol. 10, veae034 (2024).

66. Han, A. X. et al. Within-host evolutionary dynamics of seasonal and pandemic human influenza A viruses in young children. eLife 10, (2021).

67. Harrington, W. N. et al. Rapid expansion of genotype D1.1 A(H5N1) influenza viruses in wild birds across North America during the 2024 migratory season. Nat. Med. 1–5 (2026).

68. Zaraket, H. et al. Mammalian adaptation of influenza A(H7N9) virus is limited by a narrow genetic bottleneck. Nat Commun 6, (2015).

69. Sobel Leonard, A., et al. Deep Sequencing of Influenza A Virus from a Human Challenge Study Reveals a Selective Bottleneck and Only Limited Intrahost Genetic Diversification. J Virol 90, 11247–11258 (2016).

70. Boeras, D. I. et al. Role of donor genital tract HIV-1 diversity in the transmission bottleneck. Proc. Natl. Acad. Sci. U. S. A. 108, E1156–63 (2011).

71. Burggraaf, S. et al. H5N1 infection causes rapid mortality and high cytokine levels in chickens compared to ducks. Virus Res. 185, 23–31 (2014).

72. Harvey, J. A., Mullinax, J. M., Runge, M. C. & Prosser, D. J. The changing dynamics of highly pathogenic avian influenza H5N1: Next steps for management & science in North America. Biol. Conserv. 282, 110041 (2023).

73. Kovács, L., Domaföldi, G., Bertram, P.-C., Farkas, M. & Könyves, L. P. Biosecurity implications, transmission routes and modes of economically important diseases in domestic fowl and turkey. Vet. Sci. 12, 391 (2025).

74. Biswas, P. K. et al. Risk for infection with highly pathogenic avian influenza virus (H5N1) in backyard chickens, Bangladesh. Emerg. Infect. Dis. 15, 1931–1936 (2009).

75. APHIS. Final report for the 2014–2015 outbreak of highly pathogenic avian influenza (HPAI) in the United States. https://www.aphis.usda.gov/media/document/2086/file.

76. United States Department of Agriculture. Poultry ‘04: Animal and Plant Health Part I: Reference of Health and Management of Backyard/small Production Flocks in the United States. https://www.aphis.usda.gov/sites/default/files/poultry04_dr_parti.pdf (2005).

77. Guinat, C. et al. Poultry farm density and proximity drive highly pathogenic avian influenza spread. Commun. Biol. 8, 1306 (2025).

78. Hoffmann, E., Stech, J., Guan, Y., Webster, R. G. & Perez, D. R. Universal primer set for the full-length amplification of all influenza A viruses. Arch. Virol. 146, 2275–2289 (2001).

79. Bolger, A. M., Lohse, M. & Usadel, B. Trimmomatic: a flexible trimmer for Illumina sequence data. Bioinformatics 30, 2114–2120 (2014).

80. Langmead, B. & Salzberg, S. L. Fast gapped-read alignment with Bowtie 2. Nat. Methods 9, 357–359 (2012).

81. Danecek, P. et al. Twelve years of SAMtools and BCFtools. Gigascience 10, giab008 (2021).

82. Koboldt, D. C. et al. VarScan 2: somatic mutation and copy number alteration discovery in cancer by exome sequencing. Genome Res. 22, 568–576 (2012).

83. Proceedings of the 9th python in science conference. Proceedings of the Python in Science Conference Preprint at 10.25080/majora-92bf1922-012 (2010).

84. Hadfield, J. et al. Nextstrain: real-time tracking of pathogen evolution. Bioinformatics 34, 4121–4123 (2018).

85. Shepard, S. S. et al. LABEL: fast and accurate lineage assignment with assessment of H5N1 and H9N2 influenza A hemagglutinins. PLoS One 9, e86921 (2014).

86. Katoh, K., Misawa, K., Kuma, K.-I. & Miyata, T. MAFFT: a novel method for rapid multiple sequence alignment based on fast Fourier transform. Nucleic Acids Res. 30, 3059–3066 (2002).

87. Nguyen, L.-T., Schmidt, H. A., von Haeseler, A. & Minh, B. Q. IQ-TREE: a fast and effective stochastic algorithm for estimating maximum-likelihood phylogenies. Mol. Biol. Evol. 32, 268–274 (2015).

88. Sagulenko, P., Puller, V. & Neher, R. A. TreeTime: Maximum-likelihood phylodynamic analysis. Virus Evol. 4, vex042 (2018).

89. Suchard, M. A. et al. Bayesian phylogenetic and phylodynamic data integration using BEAST 1.10. Virus Evol. 4, vey016 (2018).

90. Lemey, P., Rambaut, A., Drummond, A. J. & Suchard, M. A. Bayesian phylogeography finds its roots. PLoS Comput. Biol. 5, e1000520 (2009).

91. Bielejec, F. et al. SpreaD3: Interactive visualization of spatiotemporal history and trait evolutionary processes. Mol. Biol. Evol. 33, 2167–2169 (2016).

92. Minin, V. N. & Suchard, M. A. Counting labeled transitions in continuous-time Markov models of evolution. J. Math. Biol. 56, 391–412 (2008).

93. Minin, V. N. & Suchard, M. A. Fast, accurate and simulation-free stochastic mapping. Philos. Trans. R. Soc. Lond. B Biol. Sci. 363, 3985–3995 (2008).

94. Rambaut, A., Drummond, A. J., Xie, D., Baele, G. & Suchard, M. A. Posterior summarization in Bayesian phylogenetics using tracer 1.7. Syst. Biol. 67, 901–904 (2018).

