## Supplementary material for "Within- and between-host dynamics of highly pathogenic avian influenza in domestic birds from Pennsylvania farms and live bird markets": Supp Figures 1-4; Supp tables 1-2; Supp Appendix 1

**Supplementary Figure 1.**

**Supplementary Figure 2.**

**Supplementary Figure 3.**

**Supplementary Figure 4.**

**Supplementary Table 1.**

**Supplementary Table 2.**

**Supplementary Appendix 1.**

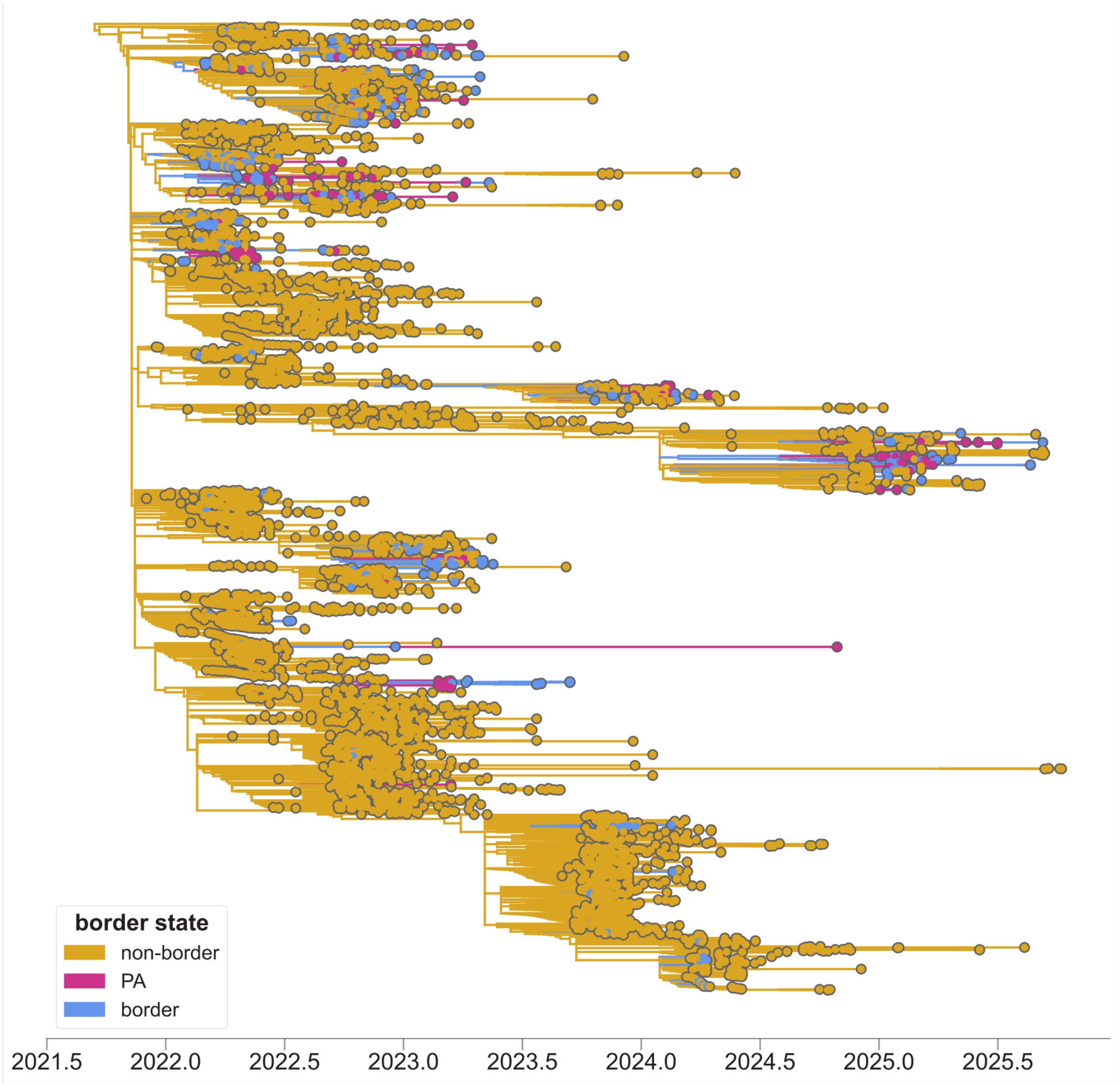


**Supplementary Figure 1.** Maximum likelihood phylogeny of 6,666 HA sequences colored by border state status. Tips are colored by domestic status. Branches are colored by the inferred trait based on the border state discrete trait analysis.

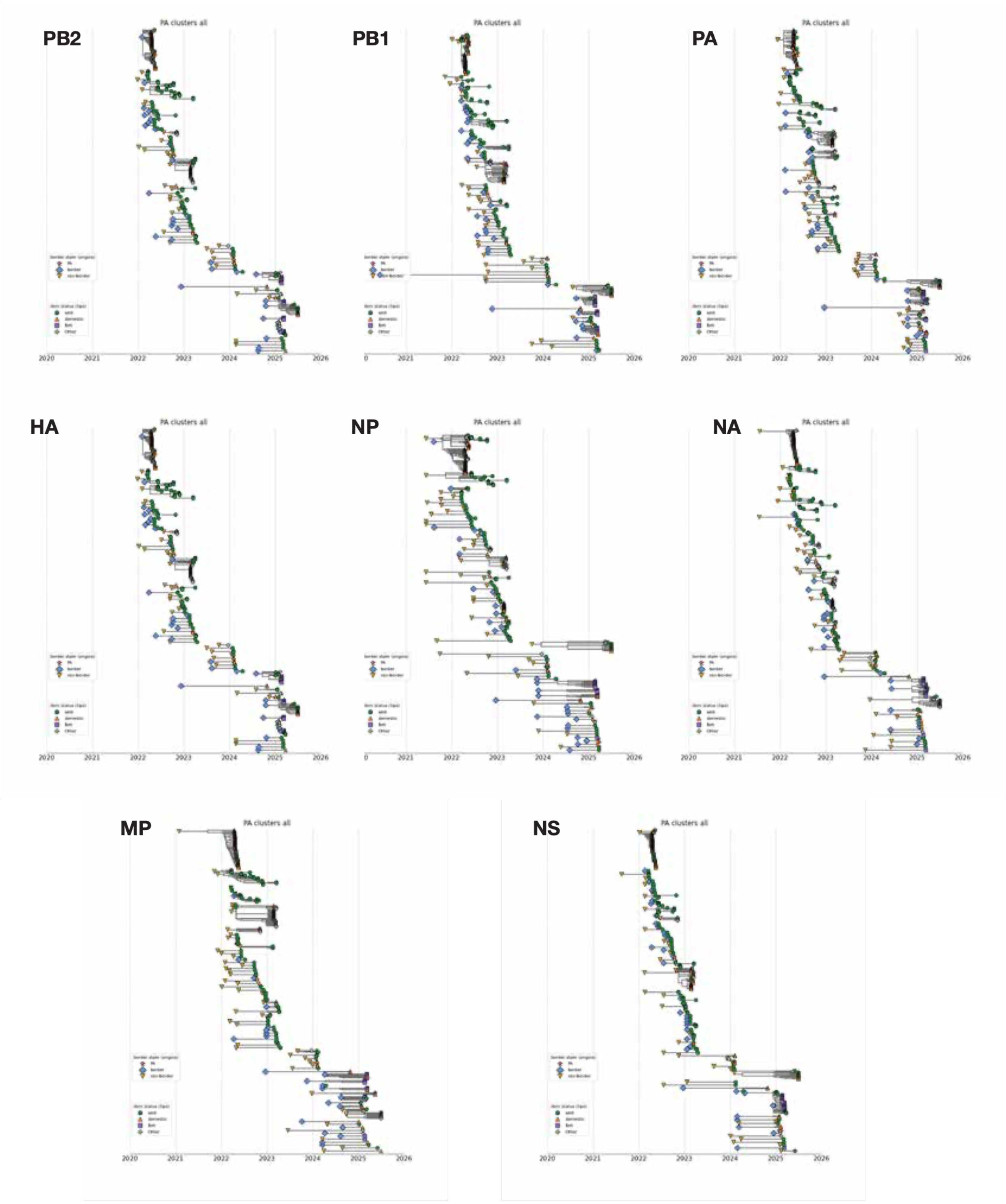


**Supplementary Figure 2.** Exploded tree view for each gene tree showing all independent Pennsylvania clusters. Tips are colored and shaped by domestic status (wild, green; domestic, yellow; lbm, purple; other,gray). Large origin tips are colored by their inferred origin location (border, blue; non-border state. yellow).

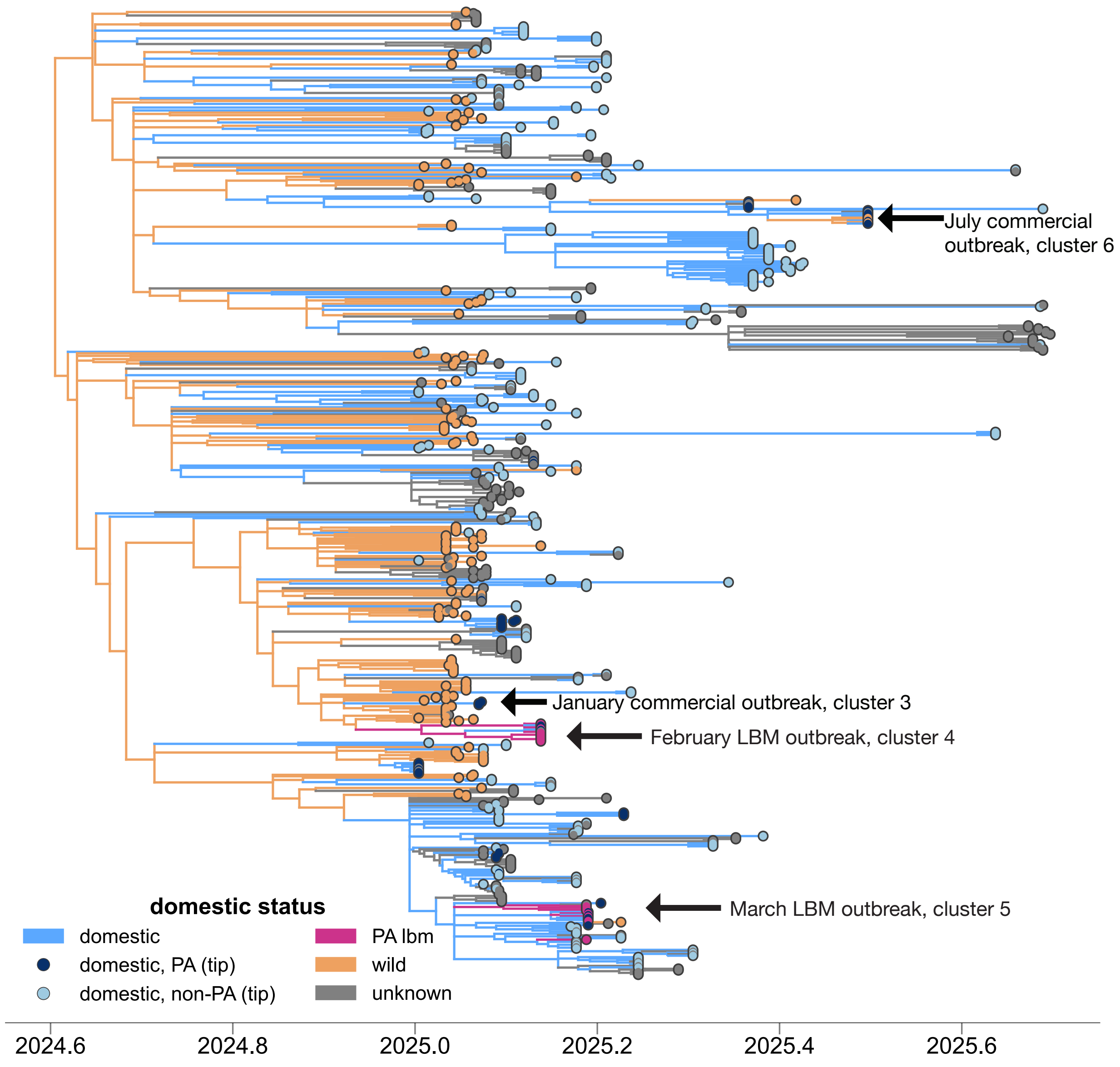


**Supplementary Figure 3.** Full-genome maximum-likelihood time resolved phylogeny of 661 D1.1 sequences. Tips are colored by domestic status. Branches are colored based on the discrete trait analysis of domestic status.

**Supplementary Figure 4.**

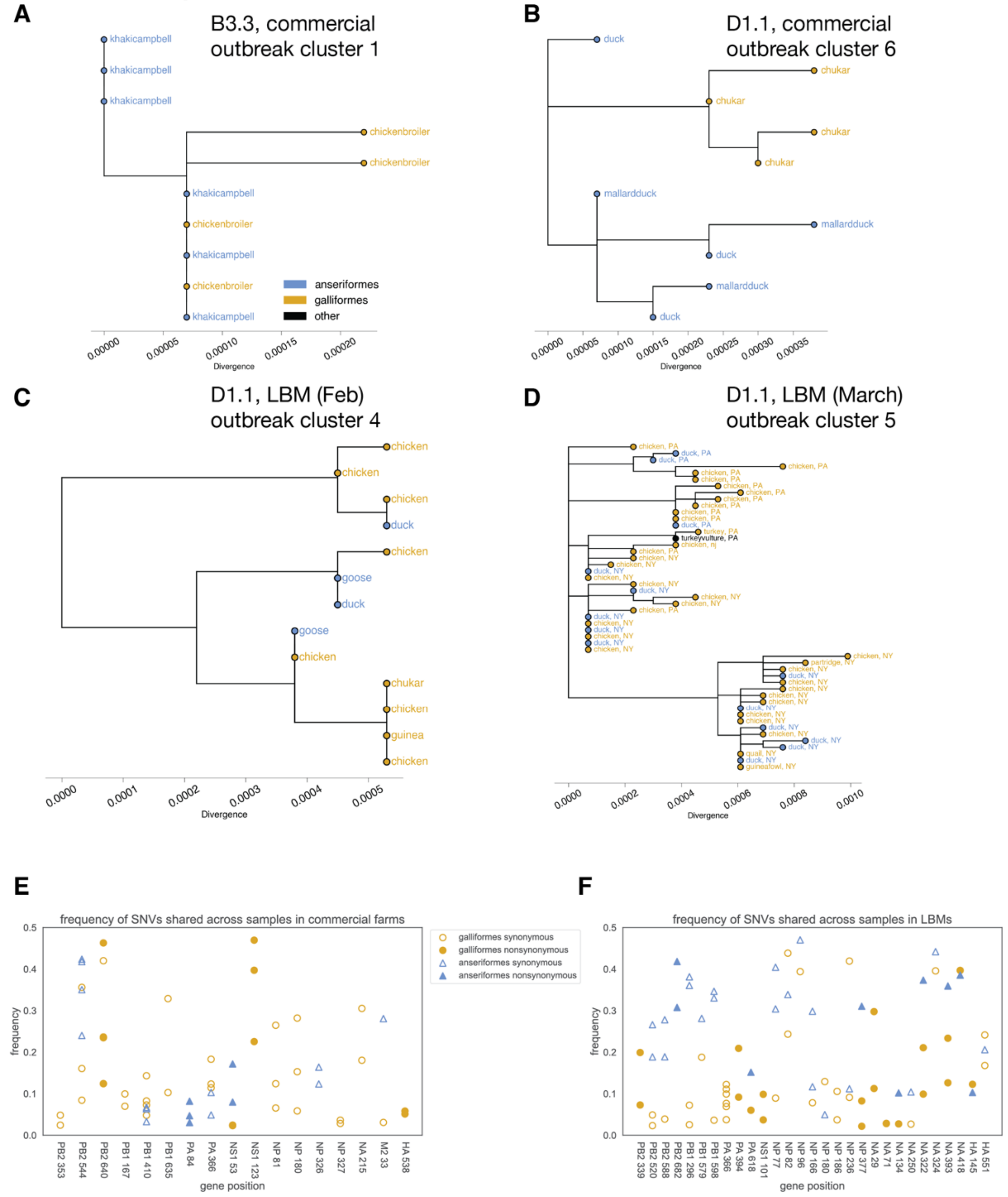


**Supplementary Figure 4.** A-D) Full-genome maximum-likelihood divergence trees of outbreak clusters with mixed species (B3.3 commercial; D1.1 commercial; D1.1 LBM February; D1.1 LBM March). Tips are colored by host order (blue, Anseriformes; yellow, Galliformes; other, black). All tips represent domestic infections with the exception of the turkey vulture tip in black in the D1.1, LBM March cluster. In the D1.1 clusters, tips are a mixture of our sequences and contextual domestic sequences from Pennsylvania. In the LBM March cluster, some sequences come from New York, which is shown by tip annotation of NY or PA. All amino acid sites that had SNVs in at least 2 samples are shown with their gene and amino acid position plotted on the x-axis in commercial farm samples (E) and LBMs (F). Open symbols indicate synonymous SNVS and closed symbols indicate nonsynonymous SNVs. Points are colored by host order.

**Supplementary Table 1. Transition rates and Bayes Factor support for D1.1 discrete trait analysis.**

| Transition | Rate (lineage/year) | ESS | Posterior Probability | Bayes Factor |
|---|---|---|---|---|
| **NY_domestic to PA_domestic** | **1.778** | **2909** | **0.995** | **484.191** |
| **NY_domestic to domestic** | **1.197** | **3052** | **0.99** | **225.770** |
| NY_domestic to wild | 0.809 | 3244 | 0.274 | 0.850 |
| PA_domestic to domestic | 0.795 | 1905 | 0.457 | 1.894 |
| PA_domestic to wild | 0.723 | 3079 | 0.415 | 1.597 |
| domestic to wild | 0.711 | 2894 | 0.443 | 1.786 |
| PA_domestic to NY_domestic | 0.737 | 1387 | 0.472 | 2.013 |
| domestic to NY_domestic | 0.635 | 3244 | 0.513 | 2.372 |
| **wild to NY_domestic** | **1.089** | **3244** | **1** | **7294.354** |
| domestic to PA_domestic | 0.711 | 3244 | 0.427 | 1.678 |
| **wild to PA_domestic** | **0.985** | **2895** | **0.999** | **3646.052** |
| **wild to domestic** | **1.817** | **3162** | **1** | **7294.354** |

**Supplementary Table 2. GISAID acknowledgement table.** Available as TSV file separately.

**Supplementary Appendix 1. GISAID EPI_SET 260701sb**

### Supplementary Appendix

All genome sequences and associated metadata supporting the findings of this study can be accessed through the persistent digital object identifier https://doi.org/10.55876/gis8.260701sb

In addition to the minted DOI, GISAID also communicates the aggregation of GISAID accession numbers (EPI_ISL_IDs) through the corresponding EPI_SET_260701sb identifier to facilitate both, the acknowledgment of all data contributors and the direct retrieval of the underlying data from GISAID used in this study.

##### Influenza Virus Segments Data Summary

| GISAID Identifier | Digital Object Identifier | Number of individual viruses | Data Collection range | Number of countries/territories |
|---|---|---|---|---|
| EPI_SET_260701sb | https://doi.org/10.55876/gis8.260701sb | 9,146 | 1905-07-14 to 2024-12-19 | 6 |
